# Merkel cell polyomavirus small tumor antigen contributes to immune evasion by interfering with type I interferon signaling

**DOI:** 10.1101/2024.06.28.601139

**Authors:** Denise Ohnezeit, Jiabin Huang, Ute Westerkamp, Veronika Brinschwitz, Claudia Schmidt, Thomas Günther, Manja Czech-Sioli, Samira Weißelberg, Tabea Schlemeyer, Jacqueline Nakel, Julia Mai, Sabrina Schreiner, Carola Schneider, Caroline C. Friedel, Hella Schwanke, Melanie M. Brinkmann, Adam Grundhoff, Nicole Fischer

## Abstract

Merkel cell polyomavirus (MCPyV) is the causative agent of the majority of Merkel cell carcinomas (MCC). The virus has limited coding capacity, with its early viral proteins, large T (LT) and small T (sT), being multifunctional and contributing to infection and transformation. A fundamental difference in early viral gene expression between infection and MCPyV-driven tumorigenesis is the expression of a truncated LT (LTtr) in the tumor. In contrast, sT is expressed in both conditions and contributes significantly to oncogenesis. Here, we identified novel functions of early viral proteins by performing genome-wide transcriptome and chromatin studies in primary human fibroblasts. Due to current limitations in infection and tumorigenesis models, we mimic these conditions by ectopically expressing sT, LT or LTtr, individually or in combination, at different time points.

In addition to its known function in cell cycle and inflammation modulation, we reveal a fundamentally new function of sT. We show that sT regulates the type I interferon (IFN) response downstream of the type I interferon receptor (IFNAR) by interfering with the interferon-stimulated gene factor 3 (ISGF3)-induced interferon-stimulated gene (ISG) response. Expression of sT leads to a reduction in the expression of interferon regulatory factor 9 (IRF9) which is a central component of the ISGF3 complex. We further show that this function of sT is conserved in BKPyV. We provide a first mechanistic understanding of which early viral proteins trigger and control the type I IFN response, which may influence MCPyV infection, persistence and, during MCC progression, regulation of the tumor microenvironment.

**Author Summary:** Merkel cell polyomavirus (MCPyV) is the only human polyomavirus that causes cancer in humans. As with all human polyomaviruses, the available infection models are limited. Thus, many processes such as the host response to infection and its regulation by the virus to establish infection and persistence are poorly understood. To better understand this interplay of viral MCPyV proteins, we performed genome-wide transcriptome and chromatin studies in primary human fibroblasts and simulated infection and tumorigenesis conditions by ectopically expressing the early viral proteins individually or in combination at different time points.

This allowed us to uncover a novel, previously undescribed function of polyomavirus sT, namely the reduction of the ISG response by affecting the ISGF3 complex, specifically by reducing IRF9 protein levels. This work sheds light on how early viral proteins influence the type I IFN response and how their interplay may affect MCPyV infection, persistence, and MCC progression.

## Introduction

Human polyomaviruses (hPyVs) show a narrow host range and high cell type restriction. Thus, current *in vitro* cell culture systems are limited, and no *in vivo* animal models exist to study the viral life cycle and/or persistence [1–5]. hPyVs persist lifelong; however, for most of them, including BKPyV, JCPyV, and Merkel cell polyomavirus (MCPyV), the sites/cell types of persistence in the natural host have not been fully revealed, and mechanisms contributing to the establishment of persistence are only poorly understood [1, 3, 5–9]. Current persistence models suggest that BKPyV establishes viral persistence with controlled viral DNA replication and limited viral progeny production only in cells mounting an innate immune response [7]. In contrast, MCPyV has been proposed to establish a different form of persistence, which is independent of viral progeny production [3, 5].

MCPyV has a small genome and thus a limited coding capacity, comprising six viral proteins with LT, sT, 57K, ALTO and one viral miRNA, MCV-miR-M1, expressed from the early region and the structural proteins VP1 and VP2 from the late region [5]. Several alternative transcripts are encoded by the early region, one of which results in the expression of an alternative to LT open reading frame (ALTO), expressed from a linear and circALTO RNA [10, 11]. ALTO plays a role in viral transcription during infection, but more complex infection models for MCPyV are needed to fully understand the function of ALTO in the viral life cycle. The early proteins small T (sT) and large T (LT) exhibit multifunctional properties and play crucial roles in infection and pathogenesis. During infection, LT initiates viral transcription and replication of viral DNA [2, 4, 12, 13]. During these processes, LT is supported by sT, although the precise role of sT in transcription and replication is not yet fully understood [2, 13]. During Merkel cell carcinoma (MCC) pathogenesis, in which sT together with a truncated form of LT (LTtr) is expressed, the viability of MCC tumor cells depends on both, sT and the tumor specific LTtr [14, 15]. However, current data suggest that sT primarily drives cellular transformation and disease progression [16, 17].

Viruses entering certain cell types, such as fibroblasts, endothelial and epithelial cells, trigger an intracellular innate immune response. This response is initiated by nucleic acid sensors called pattern recognition receptors (PRRs) that identify foreign RNA or double stranded (ds)DNA. RIG-I-like receptors (RLRs), cyclic GMP-AMP synthase (cGAS), interferon-gamma-inducible protein 16 (IFI16), and Toll-like receptors (TLRs) are prominent examples of these sensors, which recruit or activate specific adapter proteins like mitochondrial antiviral signaling protein (MAVS), stimulator of interferon genes (STING), myeloid differentiation primary response 88 (MyD88) or TIR domain-containing adaptor protein inducing Interferon-beta (TRIF) upon virus sensing. Subsequently, these proteins then recruit kinases that, in turn, activate transcription factors such as nuclear factor κB (NF-κB), activator protein 1 (AP-1), interferon regulatory factors 3 (IRF3) and IRF7. To activate the transcription of the interferon beta (IFN-β) gene (*ifnb1*), NF-κB and IRF3 or IRF7, together with an AP-1 heterodimer of activating transcription factor 2 (ATF2) and c-Jun, are necessary [18, 19]. IFN-β released from cells binds and activates the class I IFN receptor, consisting of IFNAR1 and IFNAR2. Once activated, the receptor initiates a signaling cascade that culminates in the activation and assembly of a multimeric transcription factor complex consisting of signal transducer and activator of transcription (STAT)1, STAT2, and IRF9, known as interferon-stimulated gene factor 3 (ISGF3) complex. Activation of ISGF3 and its binding to specific promoters ultimately results in the induction of hundreds of interferon-stimulated genes (ISGs) [19–21].

hPyVs are recognized by TLRs in endosomes during the entry process, such as TLR9, which recognizes CpG-unmethylated dsDNA [22, 23]. In addition, cytosolic nucleic acid sensors and specific signaling proteins, such as cGAS-STING contribute to the early recognition of a DNA virus infection [18]. The knowledge of MCPyV infection and immune regulation is highly limited with only few reports resulting mainly from overexpression studies. For instance, LT expression reduces TLR9 expression in epithelial and MCC cell lines [23]. Expression of both LT and sT in human immortalized fibroblasts increases the expression of genes regulating cell growth and cellular motility [24], a process accompanied by the induction of several inflammatory response genes, including ISGs such as oligoadenylate synthetase (OAS) 1 and IFN-stimulated exonuclease gene 20 (ISG20), the cytokines IL-1β and IL-6, and chemokines such as C-X-C motif chemokine 1 and 6 (CXCL1 and CXCL6). Moreover, sT expression, in the absence of LT, was demonstrated to inhibit NF-κB-mediated inflammatory signaling by interacting with the protein phosphatase 4 catalytic subunit (PP4C) and the NF-κB essential modulator (NEMO) [25, 26].

Previous reports observed cGAS-STING activation and induction of NF-κB signaling pathways with subsequent expression of antiviral ISGs and inflammatory cytokines in primary human skin fibroblasts infected with MCPyV at late stages of infection [27], but not during the early stages of infection [28, 29]. Also, in MCC cell lines the cGAS-STING pathway seems to be active [28, 29]. Furthermore, MCC cell lines and primary MCC tumors display an immune evasive phenotype by low Human Leukocyte Antigen class I (HLA-I) expression, which can be reversed in MCC cell lines by administration of histone deacetylase inhibitors [30–32] or reduction of sT expression [33].

In this study, we systematically analyzed transcriptional and epigenetic changes induced by the early viral gene products sT, LT and LTtr. We use overexpression analysis in primary cells and RNA-Sequencing (RNA-Seq) and chromatin immunoprecipitation (ChIP)-Seq experiments at different time points after lentiviral transduction to mimic gene expression of the early viral proteins and the resulting modulation of host gene expression during infection and pathogenesis. With this approach, we identified novel immunomodulatory properties of early viral proteins, in which sT contributes substantially to immune evasion. We show that sT acts as an antagonist by reducing immune activation induced by LT or tumor-expressed LTtr. In addition, independent of its function to reduce NF-κB signaling and proinflammatory cytokine response, sT inhibits signaling downstream of IFNAR by reducing IRF9 protein levels, thereby interfering with the ISGF3-dependent ISG response. These findings contribute to our understanding how MCPyV can successfully establish primary infection and persistence. In addition, our results are relevant for MCC pathogenesis, in which tumor cell recognition by tumor-infiltrating T cells might be inhibited by IFN-mediated immune activation in the tumor microenvironment.

## Results

### Ectopic expression of MCPyV T antigens leads to perturbations of the transcriptional profile in neonatal human dermal fibroblasts

Currently, the existing *in vitro* infection models for studying MCPyV are insufficient to fully depict the viral life cycle [4, 5, 34]. While semi-permissive replication systems can mimic some aspects, such as early gene expression, viral DNA replication, and late gene expression, they do not support significant production of infectious viral particles, viral spread and reinfection with newly formed viruses. Although an infection model based on human primary skin fibroblasts has allowed substantial progress, this model also fails to detect fully permissive infection including viral spread using standard *in vitro* cell culture techniques [27]. To test whether this model allows to interrogate gene expression changes and chromatin alterations triggered by the early viral gene products LT, LTtr and sT, we infected primary neonatal human dermal fibroblasts (nHDFs) with MCPyV particles according to the protocol from Liu and colleagues. We were able to infect approximately 5-8% of the cells, as measured by early viral protein, LT- and late protein VP1-positive cells in immunostainings after 7 days post infection (S1 Fig). However, there was no evidence of a permissive infection, such as release of new particles and re-infection, as the frequency of infected cells did not increase over time. Furthermore, the number of infected cells was too low to address host gene expression changes. Therefore, we established an overexpression protocol with lentiviral transduction, which allowed us to transduce individual T antigens (T-Ags) alone and in combination in nHDFs (Fig 1A). Thus, we could control expression levels and the time points after transduction and thereby mimic infection scenarios with early viral T-Ag expression corresponding to possible initial infection, prolonged infection, or expression patterns in tumor cells (Fig 1A-B). We used nHDFs from two independent donors to study the transcriptional and epigenetic changes induced by sT, LT, or LTtr. The LTtr protein used contains the first 257 amino acids of the protein, including the retinoblastoma protein (Rb) binding site LxCxE and, like all truncated MCC LT proteins, no longer contains a complete origin-binding domain and thus has lost the ability to support viral DNA replication.

**Fig 1.**
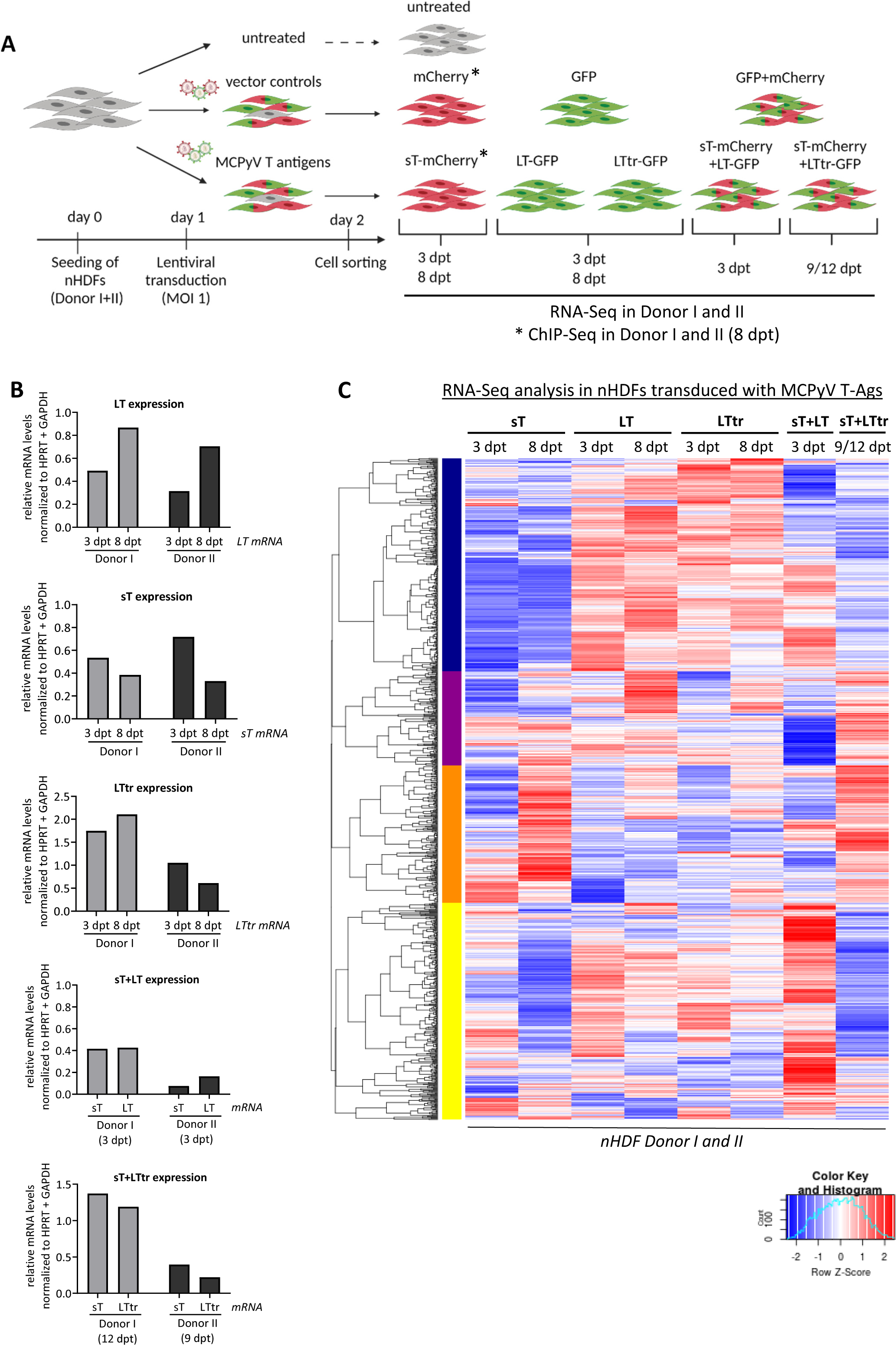
Overexpression of MCPyV T-Ags profoundly alters the host gene expression pattern in primary nHDFs. (A) Experimental outline to study transcriptional and histone modification changes induced by MCPyV T antigens. nHDFs were transduced with lentiviruses (MOI of 1) harboring MCPyV open reading frames encoding sT, LT or LTtr, in addition to GFP or mCherry. Cells were sorted at 2 dpt, followed by harvesting at 3, 8, 9 or 12 dpt for subsequent RNA-Seq and ChIP-Seq analysis. In addition to individual transductions for expression of sT, LT or LTtr, co-transductions of LT and sT or LTtr and sT were performed. Each experiment was performed once in donor I and once in donor II. (B) RT-qPCR analysis summarizing the relative mRNA levels of *LT*, *sT* or *LTtr* normalized to *HPRT* and *GAPDH* in the experiments described in Fig 1A. (C) Comparative heat map of DEGs obtained from all RNA-Seq analyses identifying four major clusters across all samples (n=2, individual donors were used as replicates). The color code refers to the row Z-scores.

Transduction efficiency and protein expression were monitored at different levels: Lentiviral transduction was performed at a multiplicity of infection (MOI) of 1 and lentiviral supernatants were titrated accordingly. mRNA expression of LT, LTtr and sT was determined at the time points by RT-qPCR and showed the expected mRNA expression of the early T constructs with some donor dependent variations in primary nHDF cells (Fig 1B). In independent experiments, the protein expression levels of the viral proteins were determined in the different donors at the different time points (S2A-B Fig). As previously described [13], we see an increased protein expression of full length LT when sT was co-expressed. In addition, we confirmed that the cDNA constructs used here do not express ALTO (S2A-B Fig).

Due to lentiviral transduction, sorting, and cell expansion, we defined the earliest time point with homogenous cell pools as immediate transcriptional changes at 3 days post transduction (dpt) and determined long-term, persistent modifications at 8 dpt. Drawing upon knowledge from other hPyVs such as BKPyV and JCPyV, which also exhibit a relatively slow infection cycle with early gene expression becoming detectable around 3-4 days post infection, we defined early time points of infection with sT and LT expression occurring 3 days post transduction.

To mimic long-term effects of early viral gene products in tumorigenesis, we focused on the effects of sT and LTtr expression at later time points. Therefore, we chose the longest possible time point of 9-12 days to consider the growth characteristics of transduced and sorted primary nHDF cells across all conditions (Fig 1). In particular, cells expressing LT alone showed limited proliferation due to genotoxic stress after early time points (donor I) and after 8 days in donor II (S2C Fig).

We performed RNA-Seq and ChIP-Seq in two independent donors of primary nHDFs for all conditions shown in Fig 1A. A comparison of total reads detected by short read sequencing is given in S1 Table. The experiments were performed once each in donors I and II at the different time points and conditions depicted, and the data from donors I and II were used as replicates in all further analyses. Transcriptional profiling of nHDFs using RNA-Seq revealed significant perturbations induced by MCPyV T-Ags at all time points, which is reflected by a large number of differentially expressed genes (DEGs, log2FC ≥1 or ≤-1 and multiple testing adjusted p-value (padj.) ≤0.05) detected in all conditions (Fig 1C, S3A, Table 1, S2 Table).

**Table 1:**
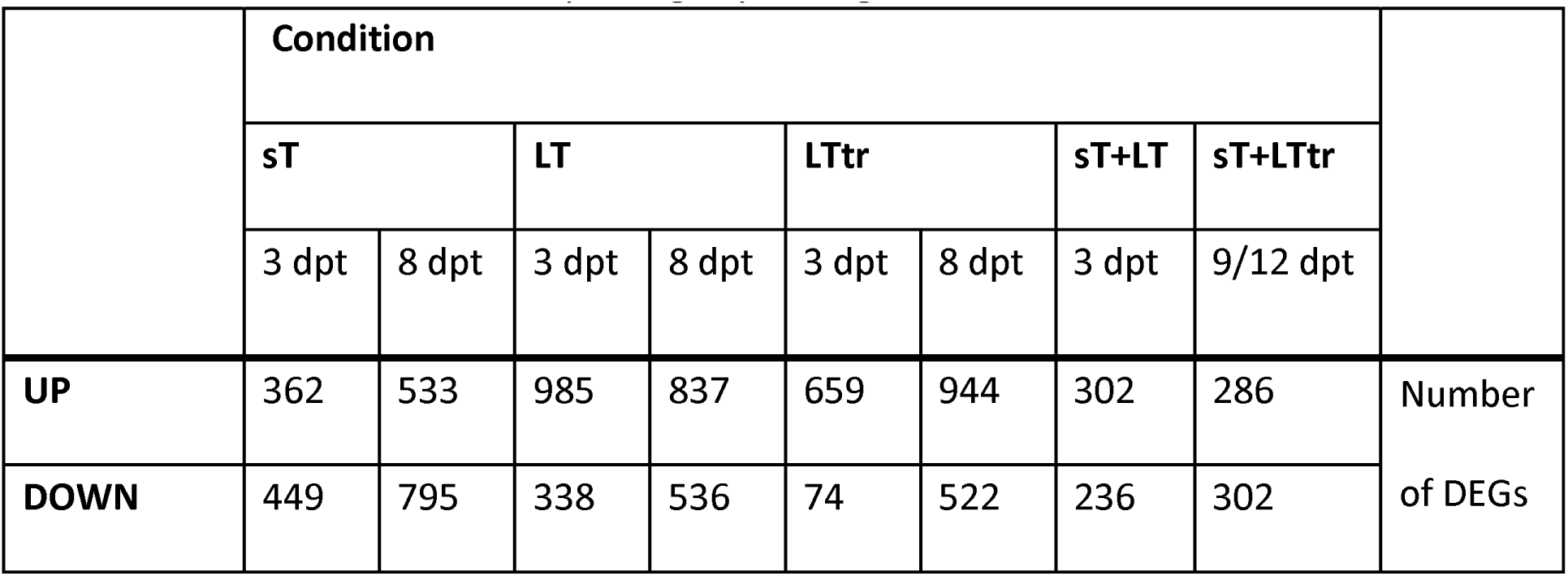
Number of DEGs in MCPyV T-Ag-expressing nHDFs. Summary of the total numbers of DEGs identified for each condition with a significant (padj. ≣0.05) upregulation (log2FC ≥1; UP) or downregulation (log2FC ≣-1; DOWN).

While principal component analysis (PCA) revealed donor-dependent clustering of samples, suggesting differential susceptibility to lentiviral transduction and T-Ag expression between donors, apparent differences in host gene expression depending on the different T-Ags were evident in the donor-specific clusters (S3B Fig).

We found several hundred differentially regulated genes in all scenarios compared to the vector control. LT- or LTtr-expressing cells displayed a trend towards more up- than downregulated genes, whereas sT-expressing cells showed similar numbers for up- and downregulated transcripts (Table 1). LT expression induced a significant upregulation of 985 at 3 dpt and 837 genes at 8 dpt. This was similar for LTtr, with 659 and 944 genes significantly upregulated at 3 or 8 dpt, respectively, while considerably fewer genes were upregulated in cells transduced with sT, sT+LT or sT+LTtr at the time points included. Interestingly, in nHDFs expressing LTtr (3 dpt), relatively few genes (n=74) were significantly downregulated (Table 1). A hierarchical clustering analysis revealed four major clusters of DEGs across all conditions (Fig 1C, S3A Fig). Based on Gene Ontology (GO) analysis, cluster I (blue) was enriched for genes involved in the innate and type I IFN response and defense response to virus (S4 Fig and S3 Table). Most of these genes were downregulated in the presence of sT but upregulated in response to the expression of LT or LTtr (Fig 1C, S3A Fig). GO analysis from genes in cluster II (magenta, Fig 1C) did not result in significant (false discovery rate (FDR) ≤0.05) terms. Interestingly, cluster III (orange, Fig 1C) was enriched for genes involved in signal transduction, inflammatory and chemokine signaling, chemotaxis, cell division, and cell adhesion (S4 Fig and S3 Table). Most of these genes were highly upregulated in cells expressing sT for 8 days or co-expressing sT with LTtr for 9-12 days (Fig 1C, S3A Fig). Meanwhile, cluster IV (yellow, Fig 1C) showed the reverse expression pattern of cluster III. This cluster was enriched for genes expressing regulators of cell adhesion, inflammatory responses, and proliferation (S4 Fig and S3 Table).

### Ectopic expression of T-Ags causes shared but also unique alterations in host gene expression patterns

The GO analysis of the DEGs found in the T-Ag-specific clusters revealed common but also distinct regulation patterns. We analyzed the data of each condition separately to investigate gene sets shared and genes specifically targeted by the individual T-Ags (Fig 2). DEGs of the different conditions are shown in S5 Fig as volcano plots and listed in S3 Table. The number of DEGs in sT-expressing cells compared to vector control are emphasized in Fig 3A. A GO enrichment analysis (for biological processes) for genes that were significantly (padj. ≤0.05) up- (log2FC ≥1) or downregulated (log2FC ≣-1) for each condition illustrates a considerable overlap in biological processes regulated by LT, LTtr and sT, e.g., cell proliferation and cell division (Fig 2 shows 10 most enriched GO terms, FDR ≤0.05 and minimal gene number of 10, see S4 Table for the complete GO term analysis, including the filtered lists shown in Fig 2 and Fig 3B). Notably, although we and others have previously found a general growth inhibitory effect by full-length LT but not LTtr expression in various cell lines, including nHDFs, the GO analysis indicated no difference in cell cycle regulation or cell division between LT and LTtr [5, 35]. However, depending on the donor and the level of LT expression, we see a significant growth inhibitory effect in cells expressing LT as previously reported. Interestingly, LTtr-expressing cells also showed a slightly reduced proliferation rate at an early stage (S2C Fig).

**Fig 2.**
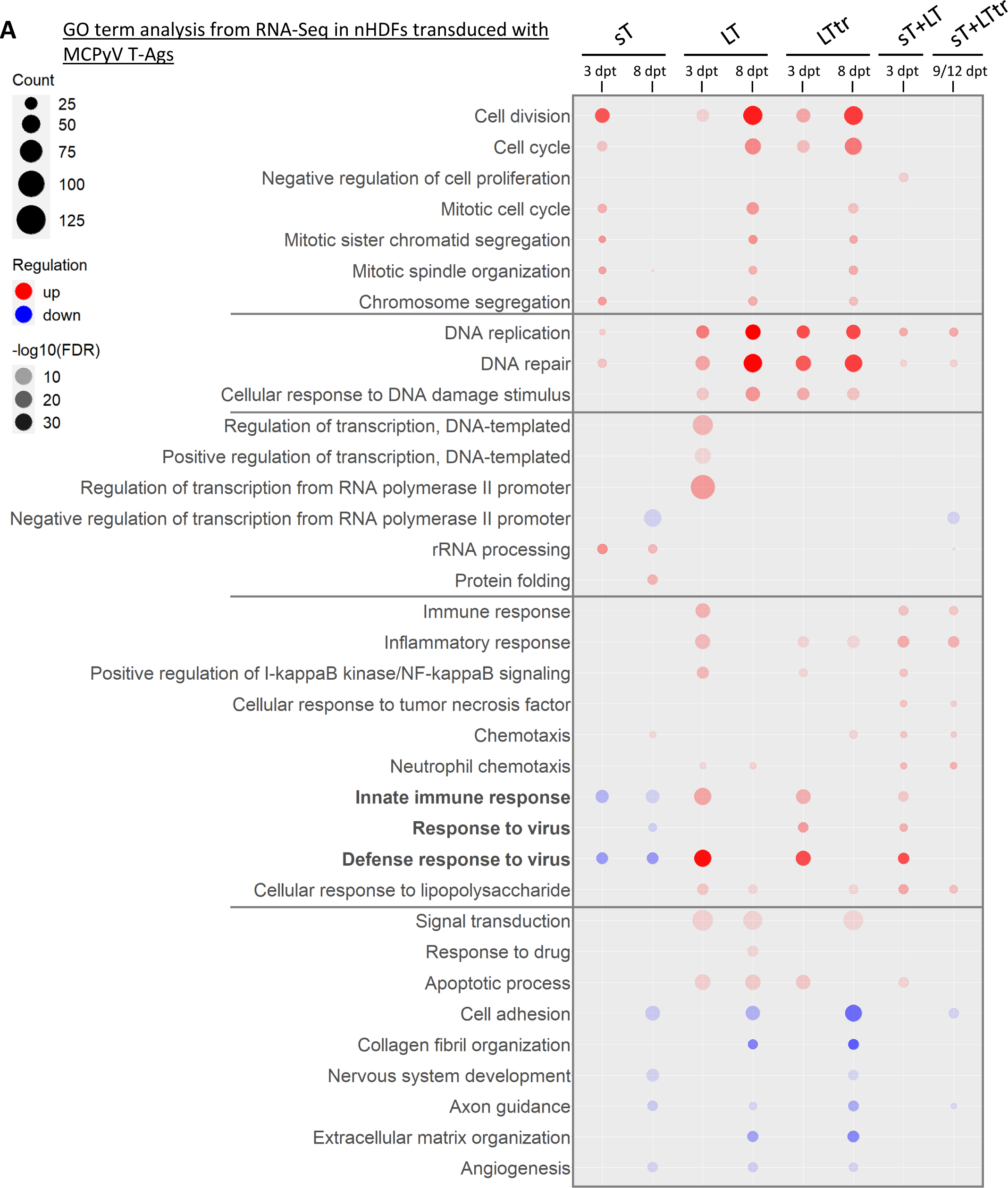
MCPyV T-Ags show an intersection of commonly deregulated and distinct host genes. GO analysis (DAVID Bioinformatics Resources) was performed using significantly (padj. ≤0.05) up- or downregulated (log2FC ≥1/≤-1) genes from the RNA-Seq analysis conducted in T-Ag-expressing nHDFs (donor I and donor II) compared to the vector controls. From all significantly enriched GO terms (FDR ≤0.05) with a minimum gene count of five, the ten terms with the highest gene counts from each condition were derived and grouped according to their biological function. The size of the icons represents the gene counts; the red color code refers to the FDR values of enriched GO terms, including upregulated genes; the blue color code indicates the FDR values of GO terms including downregulated genes. The shading represents the level of significance (FDR).

**Fig 3.**
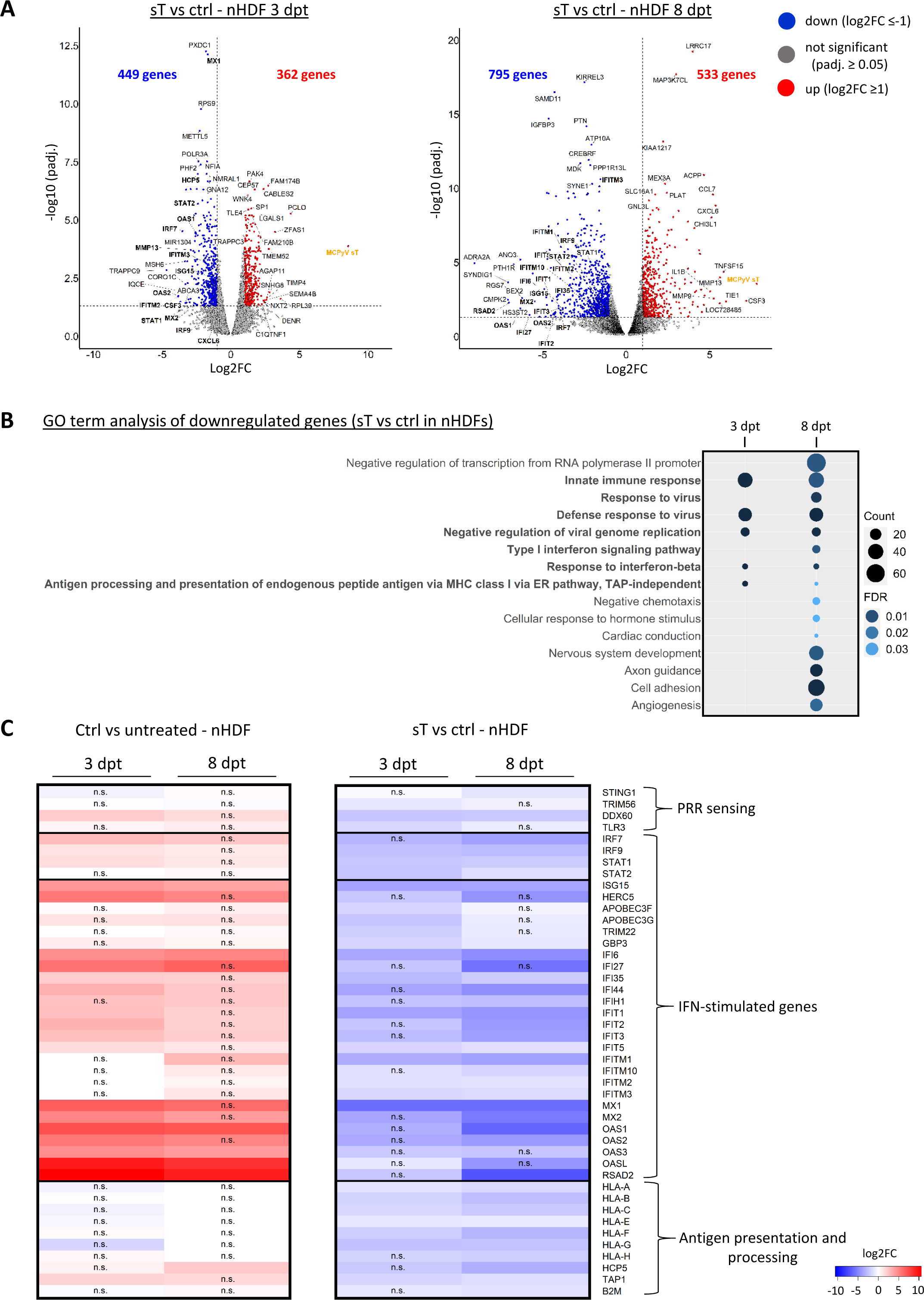
MCPyV sT suppresses the transcription of type I IFN response genes. (A) Volcano plots depicting all DEGs in nHDFs expressing sT for 3 or 8 days. RNA-Seq analysis was performed once in donor I and once in donor II, donor I and II results were used as replicates. Genes that are significantly (padj. ≤0.05) up- (log2FC ≥1) or downregulated (log2FC ≤-1) are shown as red or blue circles, respectively. MCPyV *sT* genes are shown in orange. Genes with the ten highest or lowest log2FCs and the ten most significantly differentially regulated genes are highlighted in the volcano plots. IFN-regulated genes are shown in bold. (B) GO analysis (Biological process) was performed using significantly (padj. ≤0.05) downregulated (log2FC ≤-1) genes in nHDFs expressing sT compared to the vector control. The size of the icons represents the gene counts, and the color code indicates the level of significance (FDR values). (C) Heat maps showing the changes in gene expression for a selection of genes involved in innate immunity, derived from the GO terms marked in bold in (B). A complete list of genes included in (B) is provided in the S5 Table. The left heat map shows the log2FCs of nHDFs at 3 and 8 dpt with lentiviruses expressing empty vector (ctrl), compared to non-transduced cells. The right heat map shows the log2FCs of nHDFs after 3 and 8 dpt with lentiviruses expressing sT compared to ctrl.

In addition to cell cycle regulation, GO terms including DNA replication and DNA repair were highly enriched in all conditions, with LT and LTtr showing a markedly higher impact on DNA replication than sT. Interestingly, LT and LTtr, but not sT, induced a moderate upregulation of genes responsive to DNA damage. Furthermore, the presence of any T-Ags had an impact on transcription-related processes. It is noteworthy; terms related to transcription regulation were specifically enriched only for genes that were downregulated by sT, opposed to LT-expressing cells, in which we found an enrichment of GO terms associated with transcription for upregulated genes.

Our findings align with previous studies, indicating that expression of MCPyV T-Ags led to the upregulation of a specific set of genes involved in innate immunity and inflammation [24, 36]. We observed that cells expressing sT in combination with either LT or LTtr exhibited a notable increase in mRNA expression of genes associated with chemotaxis, response to tumor necrosis factor (TNF), and NF-κB signaling. In addition, as reported previously, we found that the presence of sT affects genes associated with angiogenesis and cell adhesion [37–39]. Our study further revealed that LT and LTtr disrupt these biological processes similarly.

Intriguingly, the most striking transcriptional changes were detected in type I IFN signaling genes and innate immunity signaling pathways. Whereas LT/LTtr and sT+LT enhanced these processes at early time points (3 dpt), they had no effect at later time points. However, in contrast to the upregulation of these genes by LT/LTtr or sT+LT expression, sT expression in the absence of LT/LTtr showed the opposite effect, leading to a repression of genes belonging to the type I IFN response irrespective of the time point. Based on our finding that ectopic expression of sT repressed the type I IFN response, together with previous descriptions that MCPyV-positive tumors, to whose development and progression sT clearly contributes, are immune evasive, we focused on the effect of sT.

### MCPyV sT suppresses the transcription of type I IFN response genes

The volcano plots in Fig 3A provide a summary of all DEGs detected in sT-expressing nHDFs compared to the vector control at 3 and 8 dpt. Notably, the most upregulated genes included *CCL7, CXCL6*, and *IL1B*, which are associated with the inflammatory response. A GO analysis of all genes upregulated after sT overexpression (S6 Fig) is consistent with the previous findings that sT promotes cell division and proliferation, and interferes with transcriptional regulation, protein biogenesis, and chemotaxis [24, 37, 39]. Interestingly, compared to LT/LTtr-Ag proteins, sT expression led to a higher number of genes with reduced gene expression (449 at 3 dpt and 795 at 8 dpt) (Figs 1C, 3A, S3A Fig). A GO analysis of all genes significantly downregulated in sT-expressing cells revealed 15 GO terms with an enrichment of at least 10 genes per term and a maximum FDR of 0.05, including genes involved in negative transcriptional regulation and inhibition of viral genome replication (Fig 3B). Moreover, GO terms such as cell adhesion, angiogenesis, and development were enriched at the later time point, 8 dpt. Interestingly, type I IFN responsive genes, explicitly involved in IFN-α/β signaling, were most strongly regulated by sT (Fig 2). In addition, sT expression also resulted in the downregulation of genes associated with antigen presentation via major histocompatibility complex (MHC) class I (Fig 3B-C). A subset of genes from GO terms involved in innate immunity and type I IFN signaling (S4 Table) were selected and grouped according to their function (Fig 3C). We found that sT repressed the expression of *STING1*, *TRIM56*, *DDX60* and *TLR3*, which are involved in cytosolic DNA and RNA sensing. Further, we found a large subset of genes downregulated in the presence of sT that are classified as ISGs, according to their definition of being expressed in response to IFN [29]. We additionally found a specific cluster of genes related to MHC-I signaling, such as *HLA-A, HLA-B, HLA-E, TAP1* and *B2M* (Fig 3C).

When we compared the transcriptional response in nHDFs transduced with the lentiviral vector controls to untreated nHDFs, type I IFN response genes were significantly upregulated indicating a temporal antiviral response mediated by lentiviral transduction (Fig 3C, left panel). Antiviral response signaling is efficiently counteracted by sT at both time points with a slightly more pronounced repression of innate immune genes by sT at 8 dpt (Fig 3C, right panel). The reduced gene expression of selected gene loci for PRR sensing, ISGs and antigen presentation mediated by sT was confirmed by RT-qPCR in sT-expressing nHDFs (S7A Fig) and in MCC cell lines in which sT expression was downregulated by doxycycline-mediated induction of a sT-specific shRNA [33] (S7B Fig). Not all differences in gene expression were statistically significant, but with the exception of IRF9, the MCC cell lines showed the expected trend of decreased gene expression when sT was overexpressed and increased gene expression when sT was downregulated.

### MCPyV sT specifically represses transcription of ISGF3-dependent genes

We found that sT, in response to lentiviral transduction, consistently repressed the transcription of genes involved in innate immunity, and specifically downmodulated ISGs, which are activated in response to IRF3 and type I/II IFN signaling. Cross-referencing these genes with the interferome database [40] revealed that 16-17% of the upregulated genes and 25-35% of the downregulated genes were classified as type I/II IFN-dependent (Fig 4A, see S5 Table for the complete analysis).

**Fig 4.**
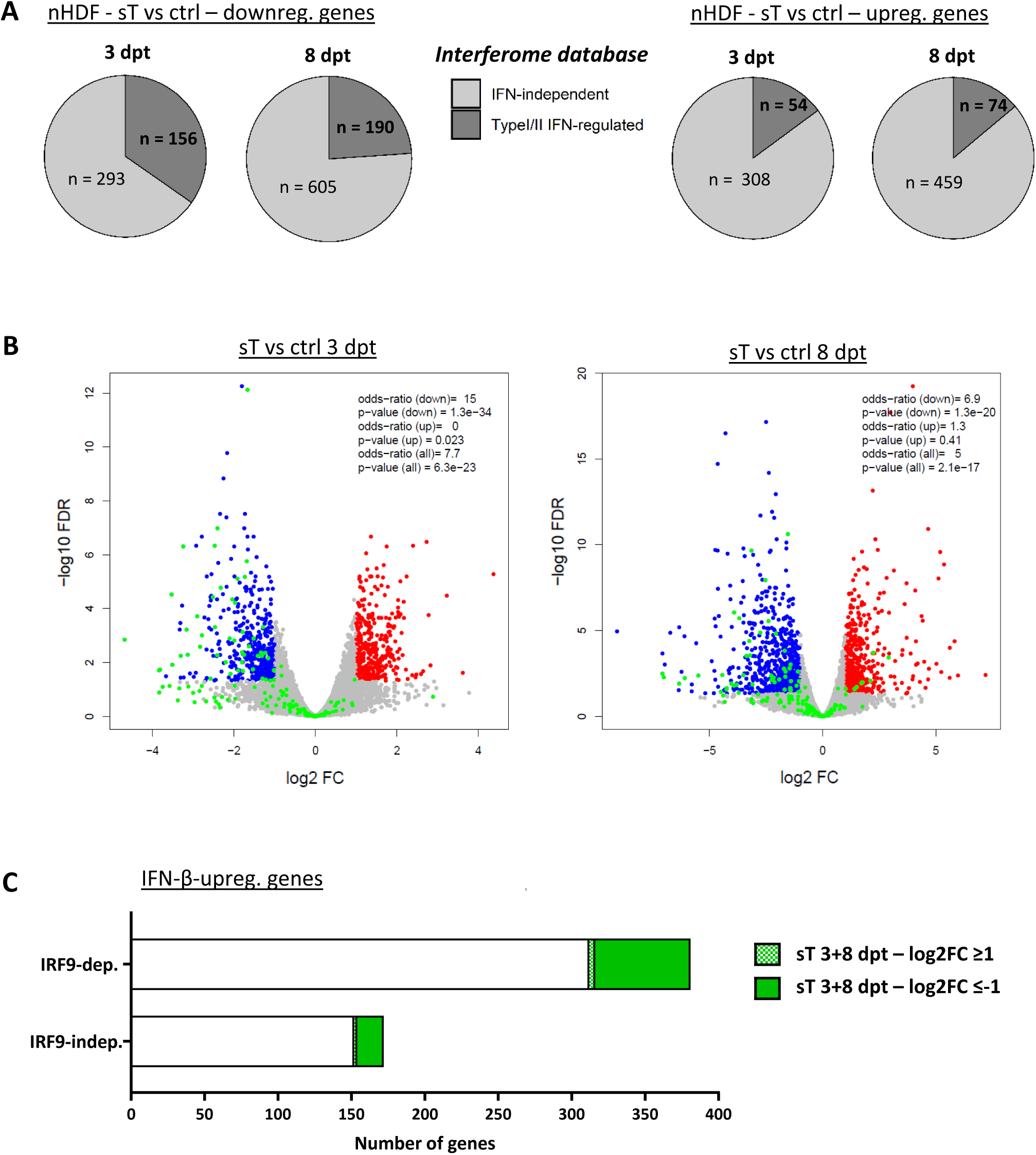
MCPyV sT affects ISGF3-dependent gene expression. (A) Comparison of genes that were downregulated or upregulated after sT expression at 3 or 8 dpt with the interferome database [40, 80]. (B) Comparison of DEGs from sT-expressing nHDFs to DEGs in WT and IRF9-KO macrophages upon IFN-β treatment from the study by Platanitis et al. [41] (NCBI GEO DataSet GSE128113). IRF9-independent (indep.) genes refer to DEGs that are regulated in the same direction in both WT and IRF9-KO cells, while IRF9-dependent (dep.) genes are DEGs in WT that are not significantly differentially expressed in the same direction in IRF9-KO cells. Statistical significance (p-values and ORs) of the overlap between DEGs from sT-expressing nHDFs and IRF9-dep. and indep. regulated genes was determined using Fisher’s exact test. Volcano plots depict all DEGs in nHDFs expressing sT for 3 or 8 days. Genes that were IRF9-dependently upregulated by IFN-β are marked in green in the volcano plot. (C) Bar plots indicating IFN-β-upregulated genes that were IRF9-dep. or IRF9-indep. are indicated on the x-axis. Genes differentially expressed in nHDFs expressing sT for 3 and 8 dpt (merged) are colored in green.

To determine whether the genes differentially regulated in the presence of sT were regulated by ISGF3, the key transcription factor complex required for ISG activation downstream of the IFNAR, we compared sT-regulated genes with previously described IRF9-dependent ISG datasets. We reanalyzed an RNA-Seq dataset from human wild type (WT) and IRF9-knockout (KO) macrophages treated with recombinant IFN-β [41, 42]. Genes were defined as IRF9-dependent if they were significantly (padj. ≤0.05) up-(log2FC ≥1) or downregulated (log2FC ≤-1) by IFN-β in WT but not IRF9-KO macrophages, and IRF9-independent up- or downregulated if they were significantly regulated in both WT and IRF9 KO macrophages. We performed statistical comparison of these gene sets with genes significantly up- and downregulated in the presence of sT at 3 and 8 dpt using Fisher’s exact test (OR = odd’s ratio determined by Fisher’s exact test, Table 2). We observed a significant (p < 10^-19^) overlap of IRF9-dependent upregulated genes with genes downregulated by sT (OR of 15 at 3 dpt; OR of 6.9 at 8 dpt, Fig 4B, Table 2). In total, of 381 genes that were IRF9-dependently upregulated by IFN-β treatment, 65 genes were also downregulated in the presence of sT at 3 and 8 dpt, while 4 genes were upregulated by sT that were dependent on IFN-β signaling (Fig 4C and S6 Table). Interestingly, among IRF9-independent genes upregulated by IFN-β, 18 genes were significantly downregulated by sT, including *STAT1* and *IRF9,* as well as *PARP9* and *DTX3L*, which are involved in ISG regulation themselves [43].

**Table 2:** Comparative analysis of sT-regulated genes with IFN-β-responsive genes [41]. RNA-Seq data from sT-expressing nHDFs was compared to an RNA-Seq data set derived from Platanitis et al. (NCBI GEO DataSet GSE128113), analyzing the responses of human WT or IRF9-KO THP1 macrophages to IFN-β treatment [41]. IRF9-independent (indep.) genes refer to DEGs regulated in both WT and IRF9-KO cells, while IRF9-dependent (dep.) genes are denoted as DEGs in WT cells that are not regulated in IRF9-KO cells in the same direction. The odd’s ratios (OR) together with the respective p-values indicate the significance of the overlap of the DEGs from nHDFs versus the IRF9-dependent and -independent DEGs derived from [41]. The OR is defined as the ratio of the odds of a sT-regulated gene being IRF9-dep./indep. to the odds of it not being IRF9-dep./indep.

| sT vs ctrl | Odd's ratio (OR) and p-value (p) |  |  |  |  |  |
| --- | --- | --- | --- | --- | --- | --- |
|  | 3 dpt<br>up + down | 3 dpt<br>up | 3 dpt<br>down | 8 dpt<br>up + down | 8 dpt<br>up | 8 dpt<br>down |
| **IRF9-indep.** |  |  |  |  |  |  |
| IFN-β-<br>downreg. | OR: 2.2<br>p: 0.19 | OR: 3.2<br>p: 0.14 | OR: 1.2<br>p: 0.57 | OR: 1.8<br>p: 0.22 | OR: 1.8<br>p: 0.33 | OR: 1.7<br>p: 0.42 |
| IFN-β-<br>upreg. | OR: 1.8<br>p: 0.066 | OR: 0.7<br>p: 1 | OR: 2.7<br>p: 0.0098 | OR: 3<br>p: 9.7e-06 | OR: 2<br>p: 0.037 | OR: 3<br>p: 0.00018 |
| **IRF9-dep.** |  |  |  |  |  |  |
| IFN-β-<br>downreg. | OR: 1.2<br>p: 0.78 | OR: 1.3<br>p: 0.67 | OR: 1<br>p: 0.72 | OR: 0.48<br>p: 0.28 | OR:<br>0.42<br>p: 0.73 | OR: 0.54<br>p: 0.58 |
| IFN-β-<br>upreg. | OR: 7.7<br>p: 6.3e-23 | OR: 0<br>p: 0.023 | OR: 15<br>p: 1.3e-34 | OR: 5<br>p: 2.1e-17 | OR: 1.3<br>p: 0.41 | OR: 6.9<br>p: 1.3e-20 |

### Transcriptional changes induced by MCPyV sT correlate with a reduction of activating histone marks

Based on previously published data on sT interfering with transcription processes, such as the recruitment of the MYC-MAX-EP400 transcription factor complex to specific gene promoters and thereby triggering activation of transcription [35] or transactivation of the lysine-specific demethylase (LSD-1) repressor complex resulting in transcriptional repression [1, 44], we further investigated the involvement of histone modifications in the promoter regulation of sT-dependent DEGs. We performed ChIP-Seq experiments in nHDF cells expressing sT or the control vector with antibodies against H3K4me3 and H3K27ac, both present in active gene promoters. Further, we used antibodies against H3K27me3 and H3K9me3 to identify changes in facultative and constitutive heterochromatin, respectively. Most changes of H3K4me3 patterns in sT-expressing cells were observed in promoter regions, while differential H3K27ac, H3K27me3 and H3K9me3 signals were detected within gene bodies or in intergenic regions (Fig 5A). While we noticed only a few changes for repressive histone modification marks, we detected a considerably higher number of genes with changes for the activating histone modifications, H3K4me3 and H3K27ac (Table 3, see S7 Table for a complete analysis of differential histone modification signal). Interestingly, increased H3K4me3 (log2FC ≥1, padj. ≤0.05) was detected for only 19 annotated genes, while a loss in H3K4me3 (log2FC ≤-1, padj. ≤0.05) was detected for 678 genes. Likewise, H3K27ac was increased for 1707 genes, but decreased for 6853 genes (Table 3).

**Fig 5.**
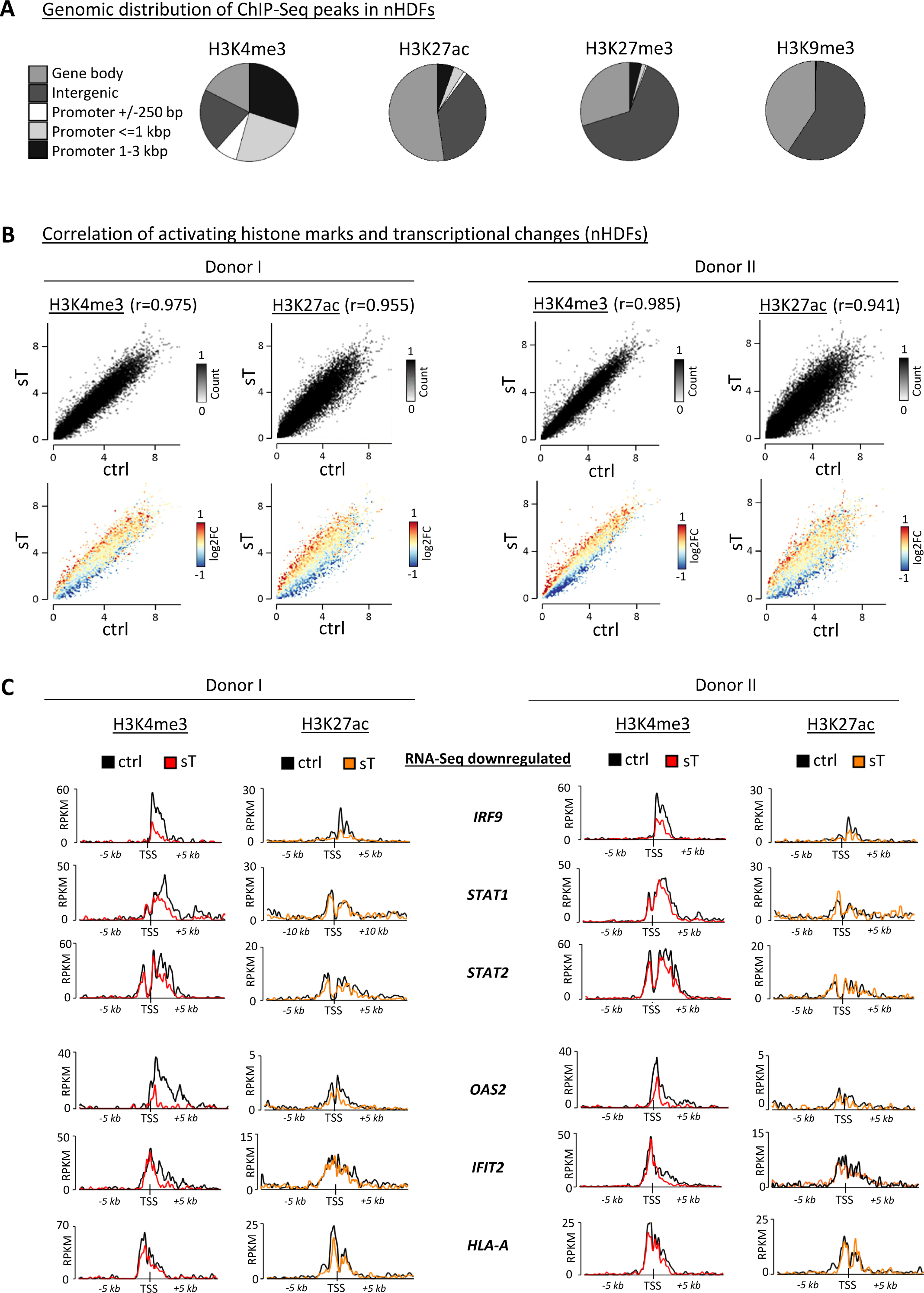
Transcriptional changes induced by MCPyV sT correlate with activating histone marks. (A) Distribution of genomic features obtained by diffReps analysis from the ChIP-Seq experiments for H3K4me3, H3K27me3, H3K27ac and H3K9me3 in primary nHDF cells overexpressing sT *vs* vector control at 8 dpt. Each ChIP-Seq experiment was performed once in donor I and once in donor II. All significant hits with a log2FC ≥0.5/ ≤-0.5 were classified according to their annotations to genomic regions. (B) Histone modification signals of activating marks, i.e., H3K4me3 and H3K27ac, were correlated with changes in gene expression, comparing sT *vs* ctrl. The color code refers to the log2FC of each gene that is plotted by its level of histone modification signal. The x and y-axes were segmented into 100 bins and regions within these bins are depicted by the counts (RPKM, reads per kb million). (C) Correlation of H3K4me3 and H3K27ac signals with gene expression data for selected genes that were downregulated in the presence of sT by RNA-Seq analysis (see Fig 3C). Average plots represent the signal (RPKM) of each histone modification within a region of 5 kb upstream and downstream of the TSS of the respective gene. Black tracks represent signals from the vector control, while tracks from sT-expressing nHDFs are shown in red for H3K4me3 and in orange for H3K27ac.

**Table 3:** diffReps hits from the ChIP-Seq analysis performed in sT-expressing nHDFs diffReps analysis for the histone modifications H3K4me3, H3K27me3, H3K27ac and H3K9me3 in primary nHDF cells overexpressing sT *vs* vector control at 8 dpt. The table summarizes the total and annotated diffReps hits with different log2FC cutoffs.

|  | H3K4me3 |  | H3K27ac |  | H3K27me3 |  | H3K9me3 |  |
| --- | --- | --- | --- | --- | --- | --- | --- | --- |
| diffReps | total | annotated | total | annotated | total | annotated | total | annotated |
| $\log_2FC \geq 0.5$ | 954 | 857 | 11238 | 6456 | 2067 | 583 | 36 | 13 |
| $\log_2FC \leq -0.5$ | 4075 | 2717 | 14335 | 9532 | 2261 | 616 | 259 | 102 |
| $\log_2FC \geq 1$ | 31 | 19 | 3458 | 1707 | 93 | 25 | 0 | 0 |
| $\log_2FC \leq -1$ | 1035 | 678 | 10298 | 6853 | 20 | 6 | 2 | 2 |

A comparison between the transcriptional activity and the enrichment for each of the histone modifications analyzed revealed a strong correlation for the activating marks H3K4me3 and H3K27ac (Fig 5B), but not for the repressive histone modification marks H3K27me3 or H3K9me3 (S8 Fig). To test whether the expression of genes within the GO terms from the RNA-Seq analysis correlated with changes in histone modification signal, we quantified the histone modification signal around the transcription start sites (TSSs), +/-5 kb of selected ISGs that were transcriptionally repressed in the presence of sT shown in Fig 3C: *IRF9, STAT1, STAT2, OAS2, IFIT2, HLA-A.* We found that their differential expression correlated with decreased levels of activating H3K4me3 and H3K27ac (Fig 5C). However, we did not detect any repressive histone modification marks near or at the promoter sites of these genes (S8 Fig). We extended our analysis to larger sets of DEGs contributing to specific GO terms (listed in the S7 Table). We did not find major changes in histone modification signal for the respective gene sets for any modification, except for H3K4me3, which was strongly reduced around TSSs of genes downregulated in the presence of sT (S9 Fig). Overall, while repressive histone modifications were not affected in the presence of sT, we found reduced H3K4me3 and H3K27ac around the TSSs of repressed genes, specifically for ISGs.

### MCPyV sT targets IRF9, a central component of ISGF3

Based on the pronounced signature of ISGF3-dependent genes among the genes downregulated by sT, we hypothesized that sT might repress ISG transcription by specifically targeting ISGF3. We confirmed a significant reduction in mRNA levels of the ISGF3 components *STAT1* and *IRF9*, as well as of the ISGs *IRF7* and *OAS2*, by RT-qPCR in sT-expressing nHDF cells compared to control cells after lentiviral transduction (Fig 6A). We also performed immunofluorescence (IF) studies and found that IRF9 expression was significantly decreased in the presence of sT compared to transduced control cells, while translocation of IRF9 to the nucleus was generally unaffected (Fig 6B, S10A Fig). We examined STAT1 phosphorylation at an earlier time point (2 dpt) to be able to detect early temporary phosphorylation events, and found a slight reduction of phospho-STAT1 (pSTAT1) in sT-expressing nHDFs compared to the vector control (Fig 6C). Similarly, IRF9 protein levels were also only slightly reduced at 2 dpt. However, at 8 dpt, we observed a significant reduction in IRF9 protein levels (Fig 6C) which we confirmed in a second donor, donor II, using the same time points described in the genome-wide RNA-Seq analyses 3 and 8 dpt. Interestingly, pSTAT2 levels were reduced in donor II after 8 days compared to the control, while we did not observe changes between sT and ctrl cells in pSTAT1 (S10B Fig). Collectively, these results indicate that sT affects the expression of ISGF3 complex genes at the transcriptional level and specifically represses the production of IRF9 at the protein level.

**Fig 6.**
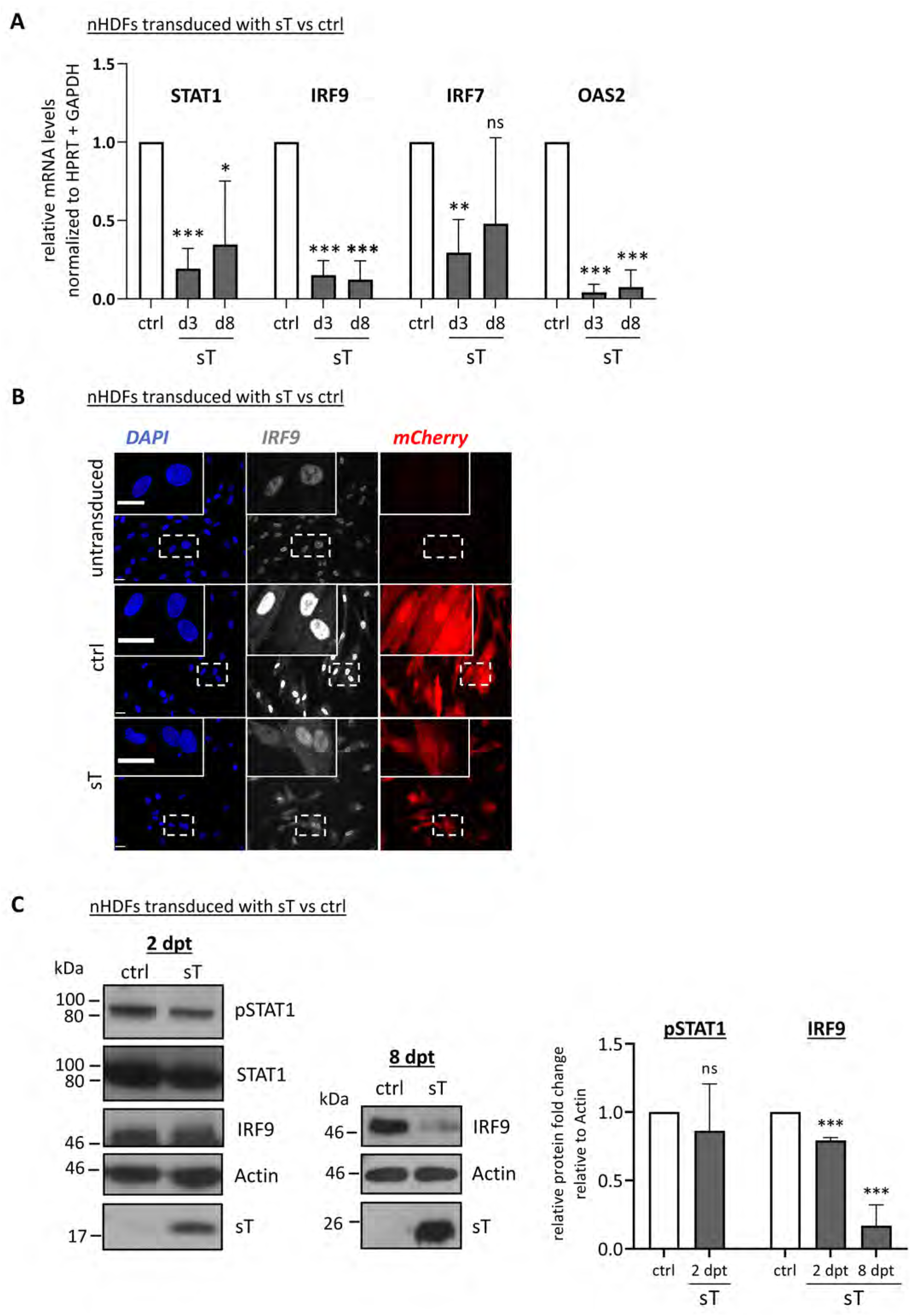
sT interferes with the signaling cascade downstream of IFNAR1/2 by targeting IRF9. (A) Confirmation of sT-induced transcriptional repression of selected ISGs: *STAT1*, *IRF9*, *IRF7* and *OAS2*. mRNA levels were measured by RT-qPCR in sT-expressing nHDFs at 3 and 8 dpt (n=3) and normalized to the mRNA levels in control cells. (B) IF staining of IRF9 in nHDFs expressing sT or ctrl vector at 3 dpt. Exemplary images are shown from non-transduced nHDFs and nHDFs expressing ctrl or sT at 3 dpt (donor II), showing DAPI-, IRF9-staining and mCherry expression, as a marker for transduction. Scale bar: 30 µm. (C) Immunoblot analysis of pSTAT1, STAT1 and IRF9 at 2 dpt (non-sorted cells) and 8 dpt (sorted cells). Representative blots are shown on the left. Quantification was performed using actin as a loading control (n=3), with normalization to ctrl cells. (A-C) Significance levels were calculated using two-tailed, unpaired *t*-test (ns = not significant, * p<0.05, ** p<0.01, *** p<0.001).

### MCPyV sT antagonizes type I IFN signaling via NF-κB and IRF9

To further investigate our findings from the RNA-Seq experiments that sT expression leads to a reduction in ISG expression, we performed luciferase-based promoter assays with the *IFNB1* promoter in HEK293 cells. When co-transfecting the *IFNB1* promoter-luciferase construct together with cGAS and STING (Fig. 7A) or RIG-I (Fig. 7B), we observed a reduction in *IFNB1* promoter activity similar to that observed with the influenza A virus (IAV) NS1 protein used as a positive control [45, 46]. However, unlike IAV NS1, we did not observe a reduction in the protein levels of cGAS, STING or RIG-I due to sT expression.

**Fig 7.**
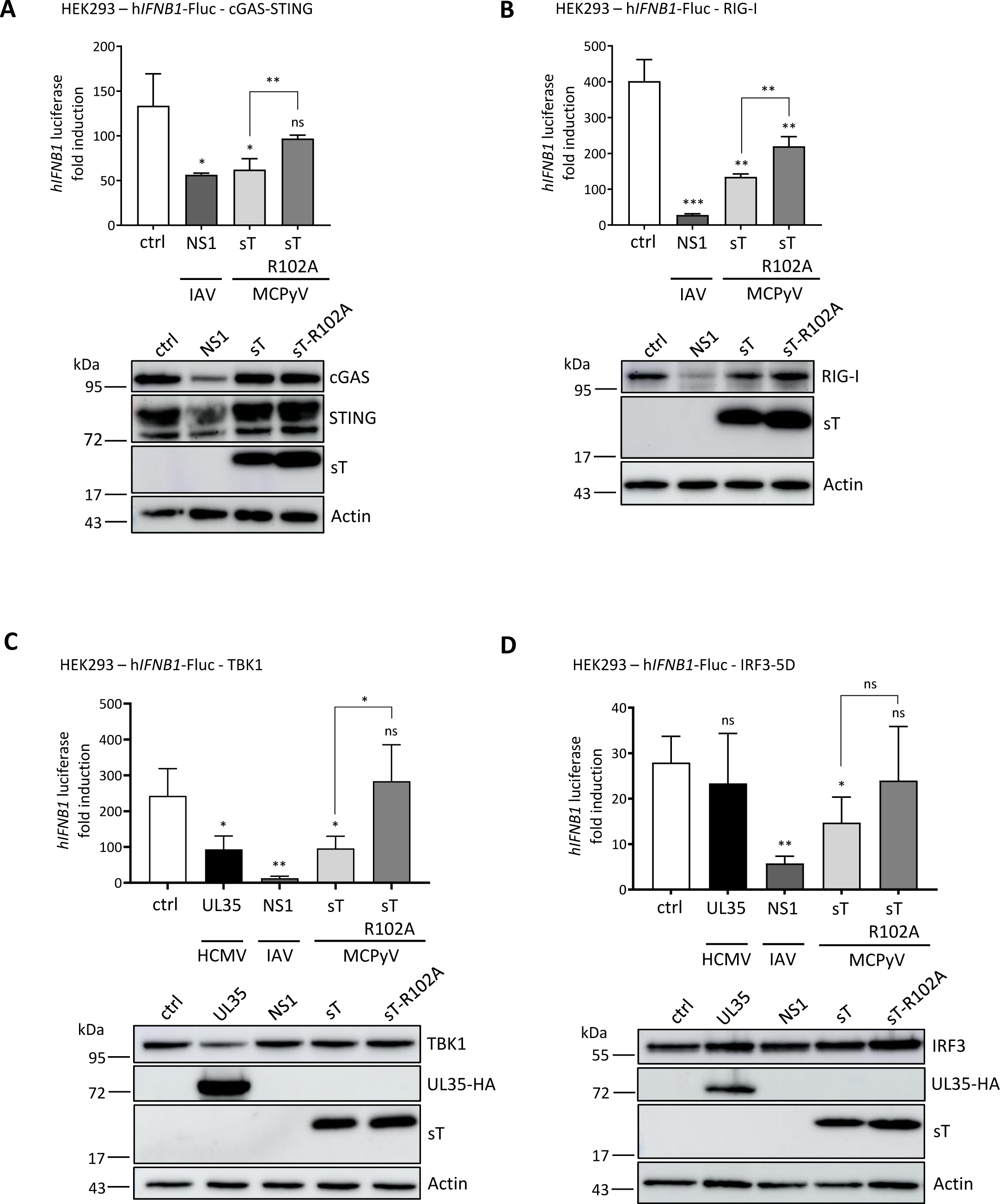
MCPyV sT downregulates transcription of *IFNB1* promoter-driven luciferase expression. (A-D) HEK293 cells were co-transfected with pRL-TK, a reporter featuring the human *IFNB1* promoter upstream of the firefly luciferase gene (*IFNB1*-FLuc), and expression plasmids for Influenza A virus (IAV) NS1, MCPyV sT, MCPyV sT-R102A or the corresponding vector control. To induce *IFNB1* promoter activity, cGAS-STING (A) RIG-I-N (B), TBK1 (C), IRF3-5D (D) or the respective controls were co-transfected and cells were lysed 20 hpt. (C-D) Expression plasmids for human cytomegalovirus (HCMV) UL35 and IAV NS1 were included as controls. Similar expression levels of viral proteins, MCPyV sT, HCMV UL35, and promoter stimuli (cGAS, STING, RIG-I, IRF3 and TBK1) was ensured by immunoblotting with protein levels of MCPyV sT analyzed using 2T2 antibody and HA antibody in the case of HCMV UL35-HA expression. Actin was used as a loading control. Representative immunoblots from one of the three independent experiments are shown below each graph. Luciferase fold induction was analyzed by normalization to *Renilla* luciferase activity, comparing stimulated *vs* unstimulated conditions. Data are shown from three independent experiments and significance levels were calculated using two-tailed, unpaired *t*-test (ns = not significant, * p<0.05, ** p<0.01, *** p<0.001).

MCPyV sT is known to target NEMO thereby inhibiting NF-κB-mediated transcription [25, 26]. To determine whether this antagonistic property contributes to the reduction in *IFNB1* promoter activity, we included a mutant of sT, sT-R102A, previously shown to rescue NF-κB-mediated inhibition of inflammatory signaling due to a decreased NF-κB essential modulator (NEMO) adaptor protein binding [25]. As expected, we find a reduced binding of sT-R102A to NEMO in cells overexpressing GFP-tagged sT/sT-R102A and FLAG-tagged NEMO (S11 Fig). Accordingly, due to impaired NF-κB activation of the sT-R102A mutant, we observe a reduced activation of *IFNB1* promoter activity in the presence of sT-R102A compared to WT sT (Fig 7A-B), with both proteins expressed at similar levels.

To further investigate whether there is a reduction in type I IFN response independent of NF-κB signaling, we examined two important signaling factors downstream of nucleic acid sensing, namely IRF3 and TBK1. While TBK1 can activate both NF-κB and IRF3, the latter is required for *IFNB1* promoter activation. When we co-transfected TBK1 with the sT proteins and the *IFNB1* promoter-luciferase construct, similar to the experiments in Fig 7A-7B, we observed that WT sT expression significantly reduced the *IFNB1* promoter activity induced by TBK1, while the sT mutant R102A showed no significant reduction (Fig 7C). Human cytomegalovirus (HCMV) UL35 and IAV NS1 served as positive controls [47].

Interestingly, when we co-expressed a constitutively active form of IRF3 (IRF3-5D), WT sT, but not sT-R102A, significantly reduced *IFNB1* promoter activity (Fig 7D). Again, UL35 and IAV NS1 served as controls with NS1 resulting in decreased *IFNB1* promoter activity while UL35 has no significant effect on IRF3-5D induced *IFNB1* promoter activation [45–47].

These results confirm previously published data that MCPyV sT affects *IFNB1* promoter activity via NEMO and inhibition of NF-κB signaling. By stimulating effectors upstream of Nf-κB signaling, the results confirm that MCPyV sT inhibits *IFNB1* promoter activity in an NF-κB-dependent manner. It is noteworthy that sT-R102A does not entirely rescue sT-mediated inhibition of *IFNB1* promoter activation.

Based on our RNA-Seq analyses in which sT counteracts lentiviral infection induced type I IFN response and our observation that sT suppresses IRF9 expression, we asked whether sT has an additional inhibitory function on the IFNAR signaling pathway. Using luciferase reporters containing the authentic promoters of the ISGs IFIT1, IFIT2 or MX1, we investigated whether sT inhibits the IFNAR signaling cascade in cells treated with IFN-β (Fig 8 and S12A-C Fig showing protein expression levels – and + IFN-β treatment).

**Fig 8.**
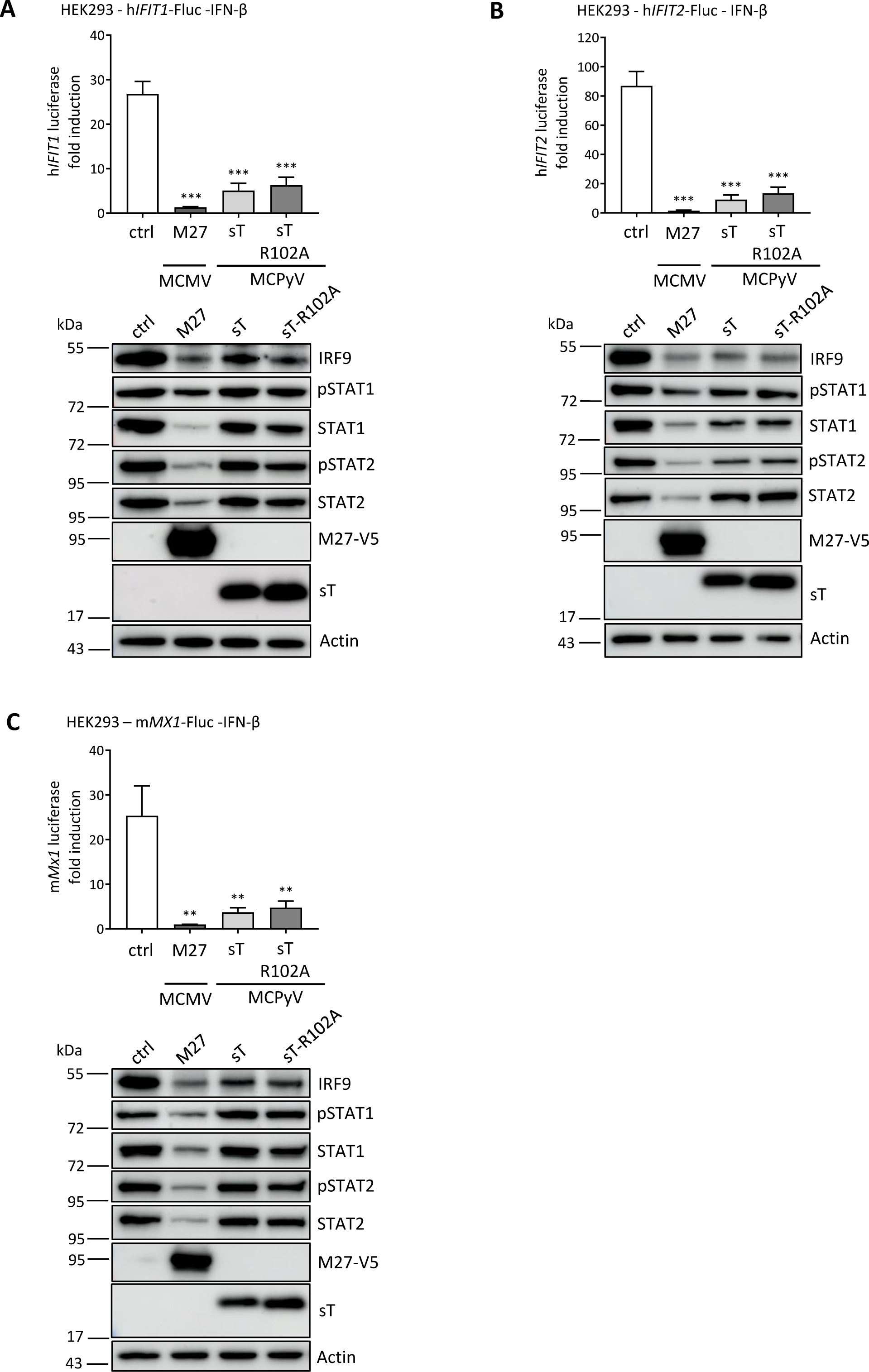
MCPyV sT downregulates transcription of *ISG* promoter-driven luciferase expression by interfering with IRF9. (A-C) HEK293 cells were co-transfected with a reporter plasmid expressing firefly luciferase under the control of the human *IFIT1* (A) or *IFIT2* (B) promoter, or the murine *MX1* promoter (C), together with pRL-TK control plasmid and expression constructs for murine CMV (MCMV) M27-V5, MCPyV sT, MCPyV sT-R102A, or the corresponding empty vector control (ctrl). 24 h post transfection (hpt), 1 ng/ml human IFN-β were added. Cells were lysed 16 h later to measure luciferase activity, comparing IFN-β-treated to unstimulated cells. Protein levels of MCPyV sT or MCMV M27-V5, pSTAT1, pSTAT2, STAT1, STAT2 and IRF9 were analyzed by immunoblotting, including actin as a loading control. Representative immunoblots of one of the three independent experiments are shown below each graph. Luciferase fold induction was analyzed by normalizing firefly to *Renilla* luciferase activity, comparing stimulated (+IFN-β) *vs* unstimulated conditions. Data are shown from three independent experiments and significance levels were calculated using two-tailed, unpaired *t*-test (ns = not significant, * p<0.05, ** p<0.01, *** p<0.001).

Similar to the control, M27 protein of murine CMV, a known antagonist of IFNAR signaling [18], overexpression of WT sT or sT-R102A mutant, both significantly impaired IFNAR-mediated ISG induction in HEK293 cells (Fig 8 A-C). The control MCMV M27 significantly reduced protein levels of all members of the ISGF3 complex, STAT1/pSTAT1, STAT2/pSTAT2 and IRF9, and particularly significantly reduced pSTAT2 levels, as previously described [48]. In contrast for sT proteins, we observed a significant reduction in IRF9 protein levels by WT sT or sT-R102A mutant expression, but only a small reduction in protein levels of STAT1/pSTAT1 and STAT2/pSTAT2, although not consistently in all three experiments (Fig 8 A-C). We provide the Western blots of the luciferase assays with and without IFN-β stimulation to verify the successful activation of the IFNAR pathway under these conditions (S12 A-C Fig).

Taken together, our results show that in addition to its known function in the type I IFN response by reducing NF-κB signaling, MCPyV sT has another independent function in the type I IFN response, specifically downstream of IFNAR activation by reducing IRF9 protein expression and thus ISG expression.

### Reduction of IRF9 protein levels due to sT expression is conserved between MCPyV and BKPyV

We asked whether this newly revealed function in the IFNAR signaling cascade, reduction of IRF9 protein levels, and reduction of ISG response is conserved for other human PyV sT proteins, particularly BKPyV sT. To this end, we repeated the luciferase assays using the *IFNB1* promoter with BKPyV sT. Since HEK293 cells do not support BKPyV sT expression, we used H1299 cells in this experiment, despite the fact that these cells would achieve a much lower stimulation in the luciferase assays. Similar to our results with MCPyV sT, we saw a reduction in *IFNB1* promoter activity when TBK1 was overexpressed together with BKPyV sT (S13A Fig). Due to the lack of commercial antibodies, we used a BKPyV sT with a V5 tag at the C-terminus. Similar to our results obtained with MCPyV sT, we also observed a reduction in *IFNB1* promoter activity when we overexpressed the constitutively active IRF3 construct, IRF3-5D (S13B Fig), although the stimulation of the cells was not sufficient in this case. Interestingly, the reduction in IFN-β-induced IFNAR activation and ISGF3-dependent promoter activity was also observed in BKPyV sT overexpressing H1299 cells (S13C Fig).

Taken together, our experiments suggest a conserved function in MCPyV and BKPyV sT proteins in reducing IRF9 protein levels and thus ISGF3 response.

### MCPyV sT counterbalances the stimulatory effect of LT and LTtron ISG transcription

Observing that MCPyV sT interfered with expression of genes involved in the host antiviral immune responses prompted us to investigate its functional consequences in infection and pathogenesis. During infection, both sT and LT are expressed and necessary for viral genome replication. Moreover, in tumor cells, the expression of sT and LTtr is crucial for cell proliferation [14, 49]. Considering both scenarios in our experimental setup, we focused on a specific group of genes enriched in the GO terms *Innate immune response*, *Response to virus*, and *Defense response to virus* (Fig 2). Most of these genes within the indicated GO terms (S5 Table) were classified as ISGs [20, 50]. We found that these genes exhibited contrasting regulatory patterns. Upon comparing their log2FCs, we confirmed that LT and LTtr, as well as the co-expression of sT+LT at 3 dpt, led to a significant upregulation of most genes. In contrast, the expression of sT alone resulted in the downregulation of these genes. Moreover, at 8-12 dpt we observed that only the expression of sT led to the repression of the selected genes (Fig 9). Notably, when inspecting the regulation of type I IFN genes, we found a significant upregulation of *IFNB1* by LT and LTtr at 3 dpt, but not by sT (Fig 9), while *IFNA1* was not significantly affected in any condition. To validate the contrasting regulation of selected ISGs (*IFI6, IRF9, ISG15, OAS2*, and *STAT1*) in the presence of sT *vs* LT or LTtr, we performed RT-qPCR on the RNA-Seq samples of the two different nHDF donors. Interestingly, we observed a slight delay in the effects of T-Ag expression, in donor I compared to donor II (S14 Fig).

**Fig 9.**
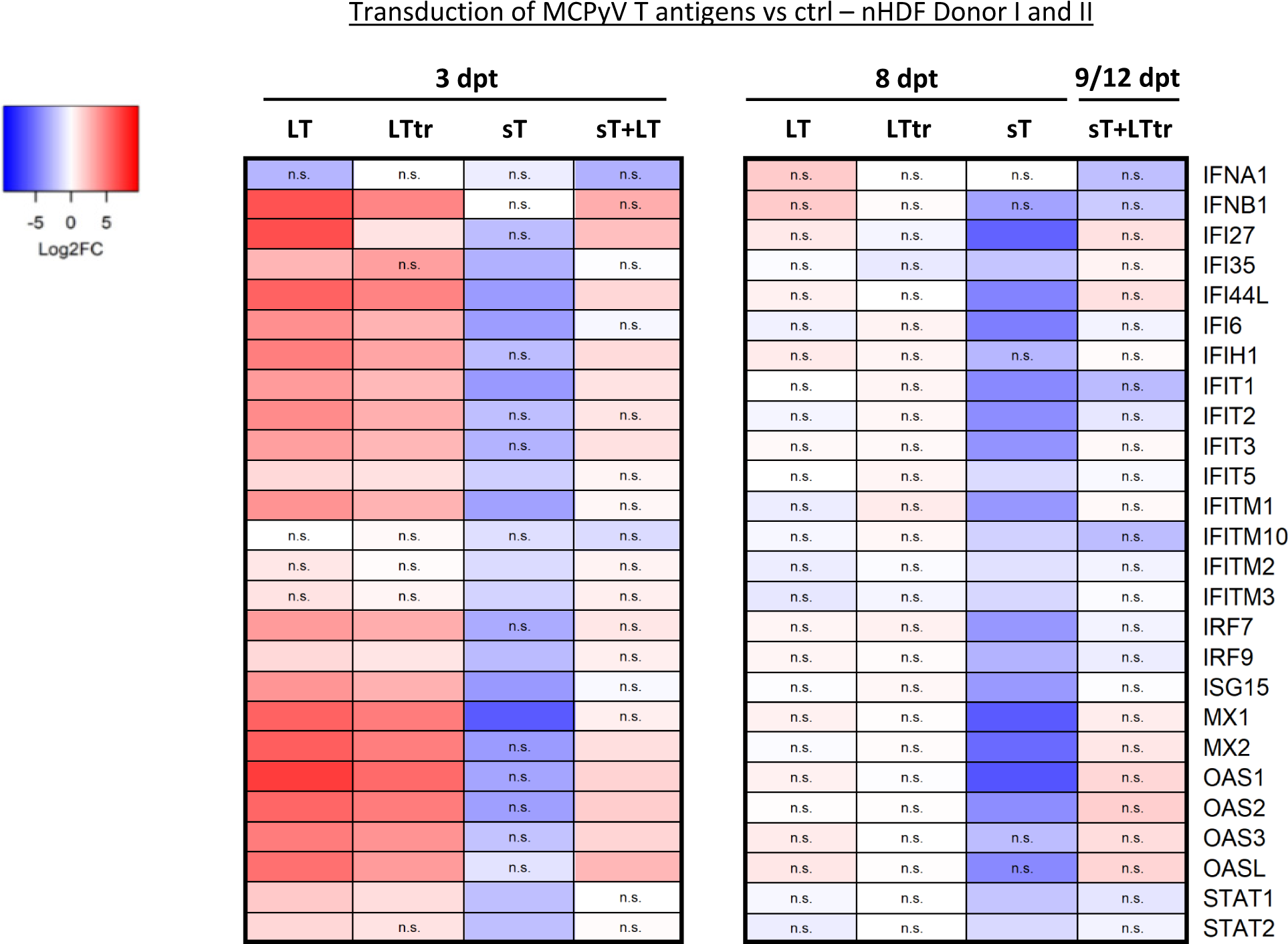
MCPyV T antigens have opposing effects on the transcription of type I IFN response genes. Heat map of selected genes involved in type I IFN signaling differentially regulated in nHDFs overexpressing MCPyV T-Ags (LT, sT, LTtr), either individually or in combination, compared to the respective lentiviral vector controls. Values are derived from the RNA-Seq analysis that was performed once in donor I and once in donor II. The color code refers to the log2FCs and n.s. is denoted for genes with a padj. value >0.05.

In summary, we found that both LT and LTtr enhanced the expression of antiviral response genes in primary fibroblasts, while sT suppressed ISG expression. Importantly, in addition to the previously described inhibition of NF-κB, we revealed IRF9 as an additional target of sT upon IFNAR1/2 signaling. By acting on multiple sites in PRR-mediated type I IFN signaling, sT is a potent inhibitor of ISG expression. We suggest that by this mechanism, sT can counterbalance transient type I IFN responses reinforced by LT in the context of infection, and by LTtr in the context of MCC progression (Fig 10).

**Fig 10.**
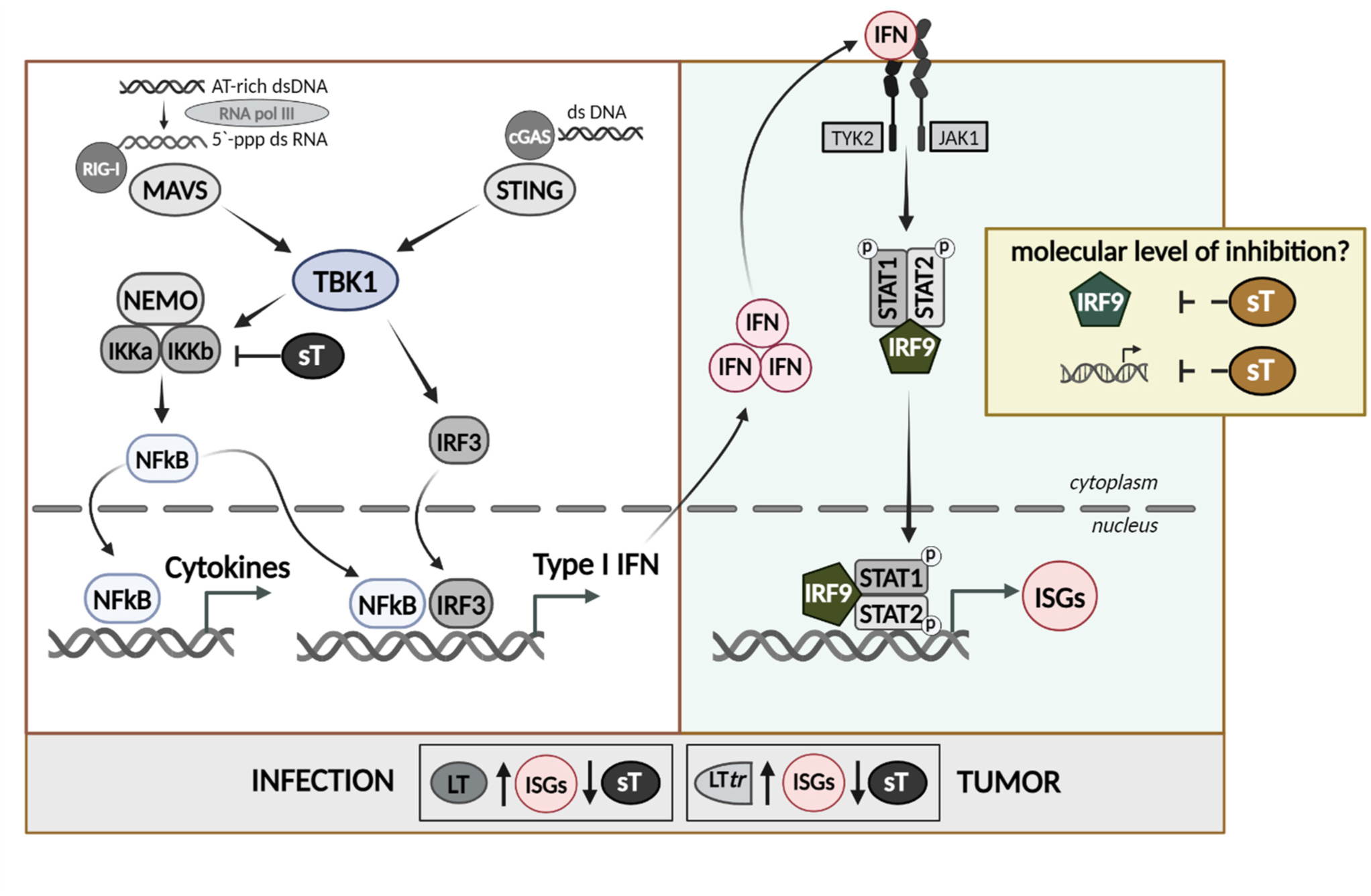
Interference of MCPyV T-Ags with PRR-mediated IFN signaling. Schematic representation of the results of this study. MCPyV sT interferes with PRR signaling by impairing NF-κB- and IRF3-mediated IFNB1 promoter activation, but additionally regulates IRF9 to repress IFNAR-mediated signaling, resulting in transcriptional silencing of a large number of ISGs. By counterbalancing the type I IFN responses elicited by LT or LTtr, sT contributes to immune evasion during infection and MCC progression.

## Discussion

MCPyV-encoded early viral proteins sT and LT work together in a tightly regulated equilibrium to facilitate viral replication during infection. Yet this interaction changes considerably in pathogenesis, partly due to selection pressure against the C-terminus of LT and expression of a truncated version of the LT protein [1, 35, 49]. To address the limitations of previous infection models, research primarily examined the effects of early viral proteins on the host response using overexpression experiments in BJ-5ta hTERT-immortalized fibroblasts [24, 51]. Similarly, investigations of the transformation properties of sT have primarily focused on understanding the mechanism of transcriptional activation and repression by sT alone, without considering the context of LT or LTtr expression [51, 52].

In this study, we conducted a thorough analysis of the transcriptional and epigenetic changes occurring in the presence of MCPyV T-Ags. Our investigation focused on primary human dermal fibroblasts, which are a relevant cell type for infection [27]. By examining the effects of early viral proteins alone as well as in combination, we have gained novel insights into the intricate interplay between the virus and viral modulation of the host immune response. We identified sT as a potent antagonist of type I IFN expression. In our experimental setup, in which we used lentiviral transduction and thereby activated the type I IFN response, we observed a sT-mediated negative modulation of the expression of several genes encoding PRR sensing molecules. Furthermore, we have identified that sT downregulates the expression of ISGs and diminishes the transcript levels of antigen presentation and antigen processing factors, including various *HLA* genes and *TAP1*. These findings further elucidate the complex interplay between the virus and the immune system important for successful infection and, importantly, for understanding viral pathogenesis.

Our results confirm and extend the scope and mechanisms of the recently published data of Krump and colleagues, who applied a complex and challenging MCPyV infection model in primary HDF cells, leading to infection of a subpopulation of the cells but not to widespread infection (see also S1 Fig) [13, 36]. By employing RT-qPCR, Krump and colleagues revealed that crucial antiviral ISGs and inflammatory cytokines exhibited increased expression levels six days post infection. They also showed that STING-TBK1-IRF3 is an important axis triggered by MCPyV infection [36]. Remarkably, no significant upregulation of antiviral genes was observed in the early stages of infection in their setup. Our results are in agreement with the study of Krump et al, consistent as that we can also observe a regulation of the STING-TBK1-IRF3 pathway.

However, our study goes beyond previous knowledge in that we show that LT is a viral factor activating inflammatory and innate immune responses while sT is the viral factor that negatively regulates innate immunity. Furthermore, our findings add on previously published data that sT mediates inhibition of NF-κB by acting downstream of receptor complexes and interacting with NEMO in the IKK complex [25, 26]. Our luciferase assays in 293 cells show, similar to the results of Griffith and colleagues, a sT-mediated reduction in NF-κB signaling and thereby reduced activation of the type I interferon response. By including the sT mutant R102A [25] that is unable to bind NEMO, we show that we have identified a novel sT function that inhibits the type I interferon response downstream of IFNAR activation independent of NF-κB inhibition. Using several authentic ISG promoters in luciferase assays after IFN-β stimulation, we show that sT inhibits ISG expression by reducing IRF9 protein expression. Furthermore, we confirm the reduction of IRF9 protein levels by sT in our lentiviral transduction experiments in primary HDF cells.

Our results suggest a balanced interaction of early viral protein expression to minimize activation of the type I IFN response: In the early stages of infection, when the expression levels of sT and LT are in equilibrium, sT effectively suppresses infection-induced PRR signaling and overrides the amplification of the response by LT. However, as infection progresses and LT induces viral DNA replication, the balance between sT and other viral factors, such as LT and newly produced viral genomes, changes. At this stage, the balance shifts in favor of factors that can trigger PRR signaling and ISG expression, rendering sT insufficient to suppress the type I IFN response effectively. Indeed, it has been demonstrated that BKPyV and JCPyV infections can elicit different ISG responses in various cell types [7, 53, 54]. For example, in certain cell types such as mouse fibroblasts or endothelial vascular cells, both BKPyV and JCPyV or BKPyV infections respectively, induce the expression of ISGs [54]. However, in primary renal proximal tubular epithelial (RPTE) cells, BKPyV fails to induce ISG expression, unlike JCPyV [7, 53]. This indicates that the ability of these polyomaviruses to trigger ISG responses may be cell-type dependent.

We have analyzed the ISG signature that is differentially regulated in the presence of sT in more detail and compared it with a dataset of IRF9 KO experiments [50]. Surprisingly, in addition to a high concordance of ISGs induced by ISGF3 or regulated in the presence of sT, we also identified a smaller subset of ISGs regulated by sT that were independent of IRF9 (Fig 4, Table 2, S6 Table). Notably, among these IRF9-independent ISGs, we identified *PARP9* and *DTX3L*. Both of these factors are known to form a complex, which as a chaperone can enhance the type I IFN response mediated by STAT1 in response to viral infection [43]. Further studies are warranted to investigate the exact mechanism by which this complex influences the IFN response triggered by MCPyV and the involved underlying molecular interactions.

In our studies on changes of histone modifications at TSSs of genes differentially regulated by sT, we have observed that the sT-mediated negative regulation of gene expression is associated with a loss of activating histone marks. Particularly, we have found a loss of H3K27ac and reduced levels of H3K4me3 after sT overexpression. These observations suggest that sT may contribute to the regulation of open and accessible promoters, rather than inducing the recruitment of transcriptionally repressive histone marks at these time points.

Consistent with previous findings on the downregulation of HLA molecules on MCC cells, our study confirms that sT plays a role in modulating the presentation of various cell surface molecules, impacting both metastasis and immune recognition [30–33]. In particular, we confirmed that the presence of sT contributes to regulation of the expression of various HLA molecules, which is also observed on MCC tumor cells, and in this way modulates their immune recognition, as we have recently shown [33]. Repression of HLA molecules in MCC tumors was also recently shown by others [30–32]. While our work demonstrates that this modulation occurs at the transcriptional level and is associated with the loss of activating histone marks, Lee and colleagues show a Polycomb-repressive complex, PRC1.1-dependent deposition of H3K27me3 at *HLA* promoters that can be reverted by inhibition of Usp7, Ubiquitin specific processing protease 7. Usp7 deubiquitinates its substrates including histone H2B thereby inducing gene silencing [32]. Differences between our observations and the results obtained in MCC cell lines may be attributable to the different cell types but even more so to the fact that MCC cell lines reflect long-term changes, some of which are also regulated by various other mechanisms.

Similar to the deregulation of HLA expression in MCC cell lines, also STING expression has been reported to be repressed in MCC cell lines and MCC tissue, similar to other cancer types [6, 29, 36]. We here provide evidence that the observed dysregulation of STING in MCPyV-positive MCC and MCPyV infection is a consequence of sT-induced repression of type I IFN-dependent genes including *STING1*. Accordingly, when we downregulated sT expression in MCC cell lines, we observed an increase in expression of *STING1* and components of the type I IFN response. The ability of sT to downregulate *STING1* expression suggests a mechanism by which MCPyV evades the host immune response thereby contributing to the progression of MCC. Thus, the modulation of *STING*1 expression by sT highlights the potential therapeutic implications for targeting sT or its associated pathways to counteract virus-positive MCC progression.

While for simian virus 40 (SV40), BKPyV, and JCPyV it has been demonstrated that LT expression induces the expression of ISGs in different cell lines [55, 56], we show for the first time a similar effect for MCPyV LT expression. Importantly, in former studies, this ISG response has been attributed specifically to the expression of LT, rather than sT, as demonstrated through cDNA overexpression experiments [54]. While in the case of SV40, increased expression of ISGs is attributed to the LT-mediated induction of DNA damage by SV40 LT [54, 55], we do not know the exact mechanism of the MCPyV LT-induced ISG response. Therefore, it will be essential to explore this further, especially since MCPyV LTtr is also capable of inducing an ISG response, and unlike LT, LTtr does not increase genotoxic stress [1, 5, 35, 49].

We additionally show for the first time that LT- or LTtr-induced gene expression of the type I IFN response is counterbalanced by sT, identifying sT as a crucial immune evasion factor. Future work in a fully permissive infection model, if once available, will be necessary to elucidate the viral pathogen associated molecular patterns (PAMPs) contributing to immune recognition and gain a comprehensive understanding of the many functions of sT in the infection cycle. Interestingly, it has recently been shown that JCPyV and BKPyV sT, but not SV40 or MCPyV sT, inhibit TRIM25-induced RIG-I activation to subvert innate immune responses, suggesting viral RNA intermediates as possible PAMPs [57]. In our experimental setup, lentiviral transduction induces the type I IFN response, which is further increased by LT/LTtr and subverted by sT. Our results are corroborated by our luciferase assays using *IFNB1* and *ISG* promoters and overexpression of sT, demonstrating that MCPyV and BKPyV sT proteins can antagonize TBK1-mediated signaling. Furthermore, we show that both MCPyV sT and BKPyV sT also act downstream of IFNAR activation in inhibiting ISGF3 dependent ISG induction in ISG promoter assays in 293 cells stimulated with IFN-β.

Limitations of our study include that due to the lack of infection models (S1 Fig), we performed overexpression experiments. However, we conducted lentiviral transduction at an MOI of 1, thus avoiding non-physiological high levels of viral protein expression and subsequent altered cellular responses. Additional constraints stem from natural differences among donors. These variations, particularly in the immune response, might affect experimental outcomes. Hence, we included two independent donors as a source of primary nHDF cells. We also did not include other cell types in our transcriptome analysis, which may exhibit distinct responses to viral proteins, and the findings thus may not be generalizable to different cell types or tissues. However, we have confirmed the results regarding the function of sT in modulation of the type I IFN response in HEK293 cells, demonstrating sT’s activity in multiple cell lines.

This study is the first to show that MCPyV sT antagonizes the type I IFN response downstream of IFNAR activation by reducing IRF9 protein levels leading to reduced ISG production. Interestingly, we show that BKPyV sT exhibits a similar immune modulatory effect downstream of IFNAR signaling with reduced IRF9 protein levels, suggesting a conserved role for PyV sT antigens in fine-tuning the host’s innate immune response to viral infection and replication products at different levels of the type I interferon response. How exactly these sT proteins antagonize the ISGF3 complex, e.g. by inhibiting IRF9 expression, inhibition of STAT1/2 phosphorylation or inducing proteasome degradation of pSTAT2 as described for other viral proteins [48, 58] will be subject of future studies.

Our findings that MCPyV sT counterbalances host responses induced by LT or, in the tumor, by LTtr, make sT an essential factor contributing to viral persistence through immune evasion but also instrumental for immune evasion of tumor cells. These findings improve our understanding of how PyVs, and MCPyV in particular, bypass innate immune responses and, importantly, offer potential targets for therapeutic approaches in MCC.

## Material and Methods

### Plasmids and Reagents

Third-generation lentiviral plasmid LeGO-MCPyV sT-iC2 (sT-IRES-mCherry) was described by us [33]. cDNA from MCPyV LT or MCC350-derived LTtr (MCC350) was cloned into LeGO-iG2 (eGFP) by restriction enzyme cloning. cDNA from MCPyV sT-co (codon-optimized) [13], or BKPyV sT was cloned into pcDNA3.1, using Gibson assembly cloning (ThermoFisher). For the generation of MCPyV sT co-R102A [25], we used site-directed mutagenesis and confirmed the correct mutation by Sanger sequencing.

For overexpression and co-immunoprecipitation experiments we used the following plasmids: pCMV-NEMO-FLAG (Addgene #11970 [59]), pMTBS-sT-YFP (Addgene #32226 [60]; pMTBS-sT-R102A-YFP, generated by site-directed mutagenesis and confirmed by sequencing.

The following plasmids were used for the luciferase-based promoter assays: pcDNA3.1 MCMV M27-V5/His (Ann Hill library) [61], HCMV UL35-HAHA [47], pcDNA3.1(-) NS1 (H1N1 influenza strain A/California/04/2009) [62] served to express viral proteins for comparison. MCPyV sT-co and BKPyV sT were cloned into pcDNA3.1(-), and C-terminally tagged with V5/His, or left untagged. pEF1α-RL (expresses renilla luciferase under the control of the EF1α promotor), pGL3basic-ISG54-FL (human *IFIT2* promoter, firefly luciferase), pGL3basic-MX1-FL (murine *MX1* promoter, firefly luciferase) and pCAGGS FLAG-RIG-I-N, expressing a constitutively active truncation mutant of RIG-I, were kindly provided by A. Pichlmair (Technical University Munich, Germany). pGL3basic-ISG56-FL (human *IFIT1* promoter, firefly luciferase) [62], pGLbasic-IFNβ-FL (human *IFNB1* promoter, firefly luciferase) [21], pcDNA3-FLAG-TBK1 [63], pIRESneo3-cGAS-GFP [64] and pEFBOS-mCherry-STING (murine mCherry-STING) [65] were described previously. pCMVBL IRF3-5D encoding constitutively active human IRF3 by containing five amino acid substitutions (S396D, S398D, S402D, S404D, S405D) was kindly provided by John Hiscott (Institut Pasteur Cenci Bolognetti Foundation, Rome, Italy).

### Cell culture

nHDFs from two different human donors were purchased from Lonza (#CC-2509). HEK293 cells (ATCC), LentiX-HEK293T cells (ATCC), H1299 cells (ATCC) and nHDFs were cultured in Dulbecco’s Modified Eagle Medium (DMEM; Gibco), supplemented with 10% fetal bovine serum (FBS; Gibco) and 1% penicillin-streptomycin (Gibco) under standard culture conditions. Patient-derived MCC cell line WaGa was cultured in Roswell Park Memorial Institute (RPMI; Gibco) medium.

H1299 cells were transfected with polyethyleneimine (PEI): 10 µg DNA were mixed with DMEM w/o supplements and PEI at a ratio of 1:10 and added to a 10 cm dish of monolayered cells. For luciferase-based promoter assays, H1299 or HEK293 cells were transfected using Fugene HD (Promega), according to the manufacturer’s protocol, at a 3:1 ratio of DNA and FuGene HD. Production of lentiviral particles was conducted in Lenti-X HEK293T cells as described [34]. nHDFs were infected as previously described [27]. ShRNA-mediated knockdown of sT in WaGa cells was performed as recently published [33].

Cellular proliferation was assessed by 3-(4,5-dimethyl-2-thiazolyl)-2,5-diphenyl-2H-tetrazolium bromide (MTT, MilliporeSigma, Burlington, MA) according to the manufacturer’s instructions. For this, nHDFs from two different donors were transduced with lentivirus expressing MCPyV sT, LT, LTtr, sT+LT, sT+LTtr or a vector control and sorted 2 days later. 1000 cells were seeded per 96-well at 3 dpt in triplicates and growth was assessed for 8 consecutive days.

### Lentiviral transduction of nHDFs

nHDFs were seeded in 6-well plates and transduced with lentiviral particles expressing MCPyV T-Ags at an MOI of 1. To increase virus uptake, polybrene (Sigma) was added (8 µg/ml) and plates were centrifuged at 37°C for 30 min at 1000 xg. Two days post transduction, mCherry- or GFP-positive nHDFs were enriched by fluorescence-activated cell sorting (FACS) and cultivated until further processing.

### GFP-TRAP experiments

Coimmunoprecipitation of ectopically expressed MCPyV-sT-YFP with NEMO-FLAG was performed by the GFP-Trap system (Chromotek) following a protocol published earlier [66]. Briefly, a total of 5lil×lil10^6^ 293 cells were transfected with 10lilμg of the C-terminal YFP-tagged WT-sT or sT-R102A or empty vector expressing YFP only, using lipofectamine 2000 (Invitrogen). Forty-eight hours posttransfection, cells were pelleted and lysed in 10× RIPA buffer according to the manufacturer’s protocol. Binding of YFP-tagged T-antigens to 20lilμl of GFP-Trap beads was carried out at 4°C for 2.5 h, with equal protein amounts used in each sample. After washing, beads were boiled for 10lilmin at 95°C in 40lilμl 2×lilSDS sample buffer, and the supernatant was analyzed by SDS-PAGE and subsequent Western blotting. The immunoreactive bands of NEMO co-precipitating with sT were detected by Image Quant 800 System (Cytiva, USA). Quantification of protein bands was performed using ImageQuant software.

### Luciferase-based promoter assays

H1299 or HEK293 cells were seeded in 96-well plates and co-transfected with 10 ng of pEF1α- RL, 100 ng of a luciferase-expressing pIFNβ-FL or pISG-FL reporter plasmid, and 100 ng of MCPyV sT, BKPyV sT, IAV NS1, human CMV UL35, or murine CMV M27 expression plasmids. Further, for each assay, a different stimulus was either co-transfected or added at a later time point, as described in the following:

### *ISG* promoter assays

pGL3basic-hIFIT1-Luc, pGL3basic-hIFIT2-Luc, or pGL3basic-mMX1-Luc were co-transfected and 24 h later, human recombinant IFN-β (1 ng/ml; PeproTech, #300-02BC) was added, or cells were left unstimulated. 16 h later, cells were lysed in 1x passive lysis buffer (PLB; Promega).

### *IFNB1* promoter assays

pGL3basic-IFNβ-Luc was co-transfected with either 100 ng of TBK1, 100 ng of IRF3-5D, 100 ng RIG-I-N or 50 ng cGAS and 50 ng STING expression plasmids. For unstimulated cells, an empty vector control was transfected instead. Cells were lysed 20 h post transfection in 1x PLB.

Luciferase activity was measured using the dual-luciferase reporter assay system (Promega), according to the manufacturer’s instructions. Luminescence was measured in a microplate reader (Tecan, Männedorf, Switzerland). Firefly luciferase signals were normalized to renilla luciferase signal. Further, the signal values from stimulated cells were divided by the values from unstimulated cells.

### Immunoblot analysis

Cell lysis was conducted using RIPA buffer (50 mM Tris-HCl, pH 7.5, 150 mM NaCl, 1 mM EGTA, 1% NP-40, 0.5% Na-deoxycholate, 0.1% SDS, 1 mM NaF, 2 mM β-glycerolphosphate, 1 mM Na3VO4, 0.4 mM PMSF, cOmplete protease inhibitor cocktail (Roche)), as described [34]. Alternatively, cells were lysed in 1x PLB for 20 min and subsequently centrifuged at 4°C for 10 min at 11,000 xg. 30-50 µg of protein were loaded per SDS gel and the following primary antibodies were used for immunoblotting: mouse mAbs, α-LT Cm2B4 (1:1000; Santa Cruz), α-sT 2T2 [67] and α-actin C4 (1:10,000; Chemicon). Further, rabbit mAbs, α-cGAS (1:1000; Cell Signaling), α-STING (1:1000; Cell Signaling), α-RIG-I (1:1000; Cell Signaling), α-IRF3 (1:1000; Cell Signaling), α-TBK1 (1:1000; Cell Signaling), α-pSTAT1 (1:1000; Cell Signaling), α-STAT1 (1:000; Cell Signaling), α-pSTAT2 (1:1000; Cell Signaling), α-STAT2 (1:1000; Cell Signaling), α-IRF9 (1:1000; Cell Signaling), α-NEMO (1:500, Cell Signaling), and α-V5 (1:1000; ThermoFisher) were used. Secondary antibodies were used as follows: Horseradish peroxidase (HRP)-conjugated mouse IgG (1:10,000; GE Healthcare) or rabbit IgG (1:3000; Cell Signaling) antibodies.

### Immunofluorescence staining, confocal and transmission electron microscopy

For IF analysis, nHDFs were grown on gelatin-coated coverslips. Cells were fixed using 4% PFA and IF staining was performed using the following primary antibodies: mouse mAb α-LT Cm2B4 (1:1000; Santa Cruz), rabbit mAbs α-VP1 (1:500; Christopher B. Buck, NCI) and α-IRF9 (1:1000; Cell Signaling). Alexa-Fluor-conjugated α-rabbit-647, α-rabbit-560, or α-mouse-488 were used as secondary antibodies, at a concentration of 1:500. Images were acquired using a confocal laser-scanning microscope (DMI 6000, Leica TCS SP5 tandem scanner, Leica, Wetzlar, Germany), using oil-immersion 40HCX or 63HCX PL CS plan-apo objectives. Image analysis was conducted in Volocity Demo Version 6.1.1 (Perkin Elmer, Waltham, MA). Quantification of IRF9 nuclear signal was performed using QuPath software [68]. From each condition, 50-150 cells were analyzed from a minimum of three coverslips per condition.

### Negative staining

Samples were adsorbed to 400 mesh carbon coated grids for 10 min. After washing with ddH2O three times, samples were stained with 1% uranyl acetate and air dried. Transmission electron micrographs were taken with a FEI Tecnai G20 equipped with an Olympus Veleta CCD camera at 80 kV.

### Quantitative PCR with reverse transcription (RT-qPCR)

For gene expression analysis, total RNA was isolated from cells using the RNeasy extraction kit (Qiagen) and DNA removal kit (Invitrogen). For RT-qPCR, 1.7 µg RNA were transcribed into cDNA using Superscript IV (Invitrogen) according to the manufacturer’s instructions. qPCR was performed using SYBR green (Thermo Fisher). A complete list of all primers and probes used in this study is given in S8 Table. Reactions were conducted in a Rotor-Gene Q-plex (Qiagen). Data were analyzed using Rotor-Gene Q Series Software (Version 2.3.1) and normalized to at least two housekeeping genes, including *TBP*, *HPRT* and *GAPDH*. Determination of MCPyV genome copies from the supernatants of infected nHDFs or BJ-5ta-immortalized fibroblasts was assessed by qPCR against VP1, including a standard derived from a MCPyV genome plasmid to calculate the genome copies.

### RNA sequencing

Isolated RNA was examined on a Bioanalyzer with the RNA Nano kit (Agilent). RNA-Seq library preparation was carried out with 1 µg of RNA from all samples that passed an RNA Integrity Number (RIN) of 0.7. mRNA enrichment was performed using the NEBNext Poly(A) Magnetic Isolation Module (NEB). RNA library preparation was conducted using the NEXTFlex Rapid Directional qRNA-Seq Kit (Bioo Scientific), according to the manufacturer’s protocol. Sequencing was conducted on an Illumina HiSeq 2500 platform (SR50). The TrimGalore (0.6.10) program (http://www.bioinformatics.babraham.ac.uk/projects/trim_galore/) was used to remove sequencing reads that contained bases with low quality scores (quality Phred score cutoff of 20) or adapters. Reads were mapped to the human genome (hg19) using the STAR 2.6.0c tool [69]. FeatureCounts (1.6.4) was utilized to calculate the read counts mapped to each gene [70]. Differential gene expression analysis was performed using DESeq2 (v1.36.0), as described [71]. GO analysis was performed with the DAVID tool [72], using DEGs with padj. ≤0.05 and log2FC ≥1/≤-1. GO terms associated with biological processes were selected with an FDR or p-value ≤0.05, and a minimum of ten genes within a GO term. All unfiltered tables from the GO term analysis are shown in S4 Table. Volcano plots, heat maps and bubble plots were generated with R Studio (4.1.0). PCA plots were created using the Python library Plotly 4.10 (https://plot.ly).

### IRF9 KO comparison

Gene read counts (annotation from GENCODE v26) for the RNA-Seq study on WT and IRF9-KO macrophages treated with IFN-β or untreated [41, 42] were downloaded from recount3 [73] (SRA ID: SRP188099, GEO ID: GSE128113). A DEG analysis comparing treated *vs* untreated cells was performed for both WT and IRF9-KO cells using DESeq2. DEGs in WT (padj. <0.05 and log2FC>1 or <-1) were considered IRF9-dependent if they were not significantly regulated in the same direction in IRF9-KO cells and IRF9-independent otherwise.

### Chromatin Immunoprecipitation sequencing

Native ChIP (nChIP) was conducted as described [74], using a starting material of 200,000 nHDFs per IP. Each IP was performed once in donor I and once in donor II. The following antibodies were used for the nChIP reactions: rabbit mAbs α-H3K4me3 (1:200; Merck Millipore), α-H3K27me3 (1:300; Cell Signaling), α-H3K9me3 (1:200; Active Motif), α-H3K27ac (1:200; abcam) and IgG (1:200; Merck Millipore). For sequencing of extracted DNA, libraries were prepared using the ChIP-Seq Library Prep Kit (Bioo Scientific) according to the manufacturer’s instructions. Samples were sequenced on an Illumina HiSeq2500, followed by trimming of the reads using the TrimGalore program (0.6.10). The trimmed reads were mapped to hg19, using the BWA tool (0.7.17-r1188) [75]. Read alignments were further processed using Samtools (1.9) [75] for tasks such as sorting, indexing and conversions. H3K4me3 and H3K27ac peak calling was conducted with MACS2 (2.1.1.20160309) [76], which is suitable for narrow peaks, while broader peaks, i.e., H3K9me3 and H3K27me3, were called using SICER (0.1.1) [77]. The diffReps tool (1.55.3) was used for the analysis of differential regions as described in [78]. Peak analyses and visualization were performed with R Studio (4.1.0) and EaSeq v.1.111 [79].

### Statistical analysis

Statistical analysis was performed using GraphPad Prism (version 9.0, GraphPad Software, San Diego,CA) and R (version 4.0.3).

## Supplementary Figures

**S1 Fig. Infection of nHDFs with MCPyV.**

(A) MCPyV total genome copies were quantified from the supernatants of infected nHDFs (n=1) or immortalized BJ-5ta fibroblasts (n=2) by qPCR targeting the VP1 gene. The arrow indicates the time point (7 days post infection) when immunofluorescence analysis (IF) was performed, shown in (B). (B) IF analysis of infected nHDF cells. LT (early viral protein, in green) and VP1 (late viral protein, in red) expression were analyzed to evaluate the infection rate. Lower panels show non-infected cells and a representative TEM of MCPyV particles used in infection.

**S2 Fig. Proliferation and protein expression of primary nHDF cells transduced with MCPyV sT, LT, LTtr.**

(A) Immunoblots from nHDFs transduced with MCPyV T antigens or the respective vector control. Transduction and sorting were performed in nHDF donor I (left) and donor II (right). Experiments shown represent independent experiments from the RNA-Seq experiments. Protein levels of MCPyV LT or LTtr were assessed using the Cm2b4 antibody, while sT was detected by the 2T2 antibody. In addition, a MCPyV ALTO-polyclonal serum was used to confirm the absence of ALTO protein production from T-Ag-specific cDNA. Actin was used as a loading control. (B) Selected samples (MCPyV LT or LTtr at 3 and 8 dpt) from the experiments described in (A) together with a positive control containing the MCPyV early region expressing ALTO were immunoblotted using the ALTO-polyclonal serum, and actin as a loading control. (C) Growth curves of nHDF cells expressing MCPyV T-Ags assessed by MTT assay in both donors, donor I and donor II. Due to technical difficulties in FACS sorting, two independent MTT assays were conducted in donor II, with the graph on the left including nHDF cells expressing sT alone or in combination with LT and LTtr, or the respective vector control, and the right graph representing nHDF cells expressing LT, LTtr or the respective vector control.

**S3 Fig. Comparison of RNA-Seq results from two nHDF donors.** (A) Comparative heat map of DEGs obtained from all RNA-Seq analyses, comparing donor I andII. The color code refers to the row Z-scores. (B) PCA plots depicting all samples that were used for RNA-Seq analysis in nHDFs from one experiment in donor I and one experiment in donor II.

**S4 Fig. GO analysis of genes differentially regulated by MCPyV T-Ags.** GO analysis was performed with DAVID Bioinformatics Resources, using all DEGs contributing to the blue, orange and yellow clusters obtained in Fig 1C. From each cluster, ten significantly (FDR ≤0.05) enriched biological processes with the highest numbers of contributing genes are depicted, as indicated behind the GO terms. The analysis from cluster “magenta” is not shown, as there were no significant GO terms detectable.

**S5 Fig. Summary of DEGs in T-Ag-expressing nHDFs.** Volcano plots summarizing all DEGs from the RNA-Seq analysis in T-Ag-expressing nHDFs from two different donors at the indicated time points. Genes that are significantly (padj. ≤0.05) up-(log2FC ≥1) or downregulated (log2FC ≤-1) are shown as red or blue dots, respectively. MCPyV *LT*, *LTtr* or *sT* are shown in orange. Genes with the ten lowest significance levels (padj.) and highest or lowest log2FCs are labeled in the volcano plots.

**S6 Fig. GO analysis of upregulated genes by MCPyV sT.** GO analysis (Biological process) was performed using significantly upregulated (log2FC ≥1) genes in nHDFs expressing sT compared to the vector control from two different donors. The size of the icons represents the gene counts, whereas the color code indicates the levels of significance (FDR values).

**S7 Fig. MCPyV sT stably represses transcription of IFN-regulated genes.** (A) RT-qPCR analysis from nHDFs (n=3) expressing sT for 3 or 8 dpt, summarizing the relative mRNA levels of *sT* and a subset of innate immune genes: *STING*, *B2M*, *TAP1*, *HLA-A* and *HLA-B*. *GAPDH* and *HPRT* were used as house-keeping genes and values were normalized to the vector control (ctrl). (B) RT-qPCR analysis summarizing the relative mRNA levels of *sT*, *IRF9*, *ISG15* and *STING* in shRNA-inducible WaGa cells (n=4) treated with Doxycycline (Dox) to induce a knockdown of endogenous sT, or DMSO. *GAPDH* or *HPRT* were used as house-keeping genes and values were normalized to DMSO-treated cells.

**S8 Fig. Transcriptional changes induced by MCPyV sT do not correlate with repressive histone marks.** (A) Signals of repressive histone marks, i.e., H3K27me3 and H3K9me3, were correlated with changes in gene expression, comparing sT *vs* ctrl in nHDF donor I and II. The color code refers to the log2FC of each gene that is plotted by its level of histone modification signal. The x and y-axes were segmented into 100 bins and regions within these bins are depicted by the counts (RPKM). (B) Correlation of H3K27me3 and H3K9me3 signals from nHDFs with gene expression data for selected ISGs downregulated by sT. Average plots represent the signal (RPKM) of each histone modification within a region of 5 kb upstream and downstream of the TSS of DEGs within the indicated GO terms.

**S9 Fig. Correlation of transcriptional and histone modification changes for specific gene sets.** Correlation of histone modification signals (H3K4me3, H3K27ac, H3K27me3 and H3K9me3) with gene expression data for selected GO terms: *Innate immunity*, *Transcriptional regulation*, *Cell adhesion*, *Cell cycle and division*, and *Protein biogenesis* . Average plots represent the signal (RPKM) of each histone modification within a region of 5 kb upstream and downstream of the TSS of DEGs within the indicated GO terms. Black tracks refer to signals from nHDFs expressing the vector control, while colored tracks represent signals from sT-expressing cells (red: H3K4me3, orange: H3K27ac, light blue: H3K27me3, dark blue: H3K9me3). Data are derived from ChIP-Seq experiments that were conducted once in donor I and once in donor II. Individual donors were used as replicates.

**S10 Fig. MCPyV sT affects ISGF3 protein expression.** (A) Quantification of immunofluorescence images from Figure 6B. Quantification was performed only on the IRF9 nuclear signal of 50-150 cells of each condition. Significance levels were calculated using two-tailed, unpaired *t*-tests (ns = not significant, * p<0.05, ** p<0.01, *** p<0.001).(B) Immunoblot analysis from nHDF donor II transduced with MCPyV sT or vector control, sorted at 2 dpt, and harvested at 3 and 8 dpt. Protein levels of IRF9, pSTAT1, STAT1, pSTAT2, STAT2 were determined by using the respective antibodies. MCPyV sT protein expression was detected using the 2T2 antibody. Actin was used as a loading control.

**S11 Fig. MCPyV sT-R102A mutant shows reduced co-precipitation with NEMO** (A) HEK293 cells were co-transfected with either a control plasmid only expressing YFP, expressing WT sT-YFP or sT R102A-YFP, in the presence of a Flag control plasmid or a plasmid expressing Flag-NEMO. After 48 h, cell lysates were incubated with GFP-Trap®Agarose beads. Bound protein was immunoblotted with sT-, NEMO-, or actin-specific antibodies. (B) Quantification of the NEMO-specific antibody signals from lane 4 and 6 (NEMO co-precipitation with WT sT-YFP and sT R102A-GFP) normalized to the input signal.

**S12 Fig. MCPyV sT reduces IRF9 protein levels under IFN-β-stimulated conditions.** (A-C) Representative immunoblots from luciferase experiments shown in Fig 8A (A), Fig 8B (B) and Fig 8C (C), indicating protein levels of IRF9, pSTAT1, pSTAT2, STAT1, STAT2, MCMV M27-V5, MCPyV sT, MCPyV sT-R102A, MCPyV sT-V5 and actin in the presence or absence of stimulation with 1 ng/ml IFN-β. Immunoblots show protein expression levels without (lanes 1-5) or with IFN-β treatment (lanes 6-10).

**S13 Fig. BKPyV sT interferes with PRR-mediated *IFNB1* and IFNAR1/2-dependent *ISG* promoter induction** (A-C) H1299 cells were co-transfected with a reporter plasmid expressing firefly luciferase under the control of the human *IFNB1* promoter (A and B) or the human *IFIT2* promoter (C), together with pRL-TK control plasmid and expression constructs for HCMV-UL35-HA, IAV NS1, MCMV M27, MCPyV sT, BKPyV sT, BKPyV-sT-V5, or the corresponding empty vector control (ctrl). Promoter activity was ensured by co-transfection of TBK1 (A) or IRF3-D (B), or by the addition of 1 ng/ml human IFN-β 24 h post transfection (C) (A-C) Protein levels of HCMV UL35-HA, MCMV M27-V5, MCPyV sT, BKPyV-sT-V5 or, pSTAT1, pSTAT2, STAT1, STAT2 and IRF9 were analyzed by immunoblotting including actin as a loading control. Luciferase fold induction was analyzed by normalizing firefly to *Renilla* luciferase activity, comparing stimulated *vs* unstimulated conditions. Data are shown from three (A-B) or 7 (C) independent experiments and significance levels were calculated using two-tailed, unpaired *t*-test (ns = not significant, * p<0.05, ** p<0.01, *** p<0.001).

**S14 Fig. MCPyV T antigens have opposing effects on the transcription of type I IFN response genes.** (A) RT-qPCR was performed from the sequenced samples described in Fig 9 for a subset of ISGs (*IFI6*, *IRF9*, *ISG15*, *OAS2*, *STAT1*).

## Supplementary Tables

**S1 Table.** Overview of read counts for each RNA-Seq and ChIP-Seq experiment.

**S2 Table.** Tabular listing of all differentially expressed genes across all different RNA-Seq experiments.

**S3 Table**. List of genes and gene ontology results of the genes listed in the color-coded clusters of the heat map Fig. 1

**S4 Table.** Complete lists of differentially expressed genes contributing to the GO Analysis shown in Figure 2.

**S5 Table.** List of Interferon stimulated genes shown in Figure 2 and 3B.

**S6 Table.** List of ISGs that are IRF9-dependently or independently regulated by sT.

**S7 Table.** Overview of the complete analysis of differential histone modification signals.

**S8 Table**. List of all primers and probes used.

## Acknowledgements

This project was funded by the Deutsche Forschungsgemeinschaft (DFG, German Research Foundation) in the framework of the Research Unit FOR5200 DEEP-DV (443644894) projects FI 782/7-1 (N.F.), GR 3318/5-1 (A.G.), SCHR 1479/5-1 (S.S.), BR 3432/7-1 (M.M.B), FR 2938/11-1 (C.C.F.), and the Helmholtz Association (W2/W3-090) (M.M.B.). The funders had no role in study design, data collection and analysis, decision to publish, or preparation of the manuscript.

We are grateful for the technical support provided by the UKE Microscopy Imaging facility (umif) and the UKE and LIV FACS facilities.

## Data availability

Sequencing data are accessible in the public repository GEO, accession number GSE230760.

## Contributions

Conceived and designed the experiments: DO NF AG MMB HS SS TG. Performed the experiments: DO JM VB HS JH UW TS SW MCZ JN CS. Analyzed the data: DO NF CCF AG TG SS MMB. Wrote the paper: DO NF.

## Notes

### Competing Interest Statement

The authors have declared no competing interest.

