## Supplementary figures and images for "Merkel cell polyomavirus small tumor antigen contributes to immune evasion by interfering with type I interferon signaling"

### supplementary Figure S1

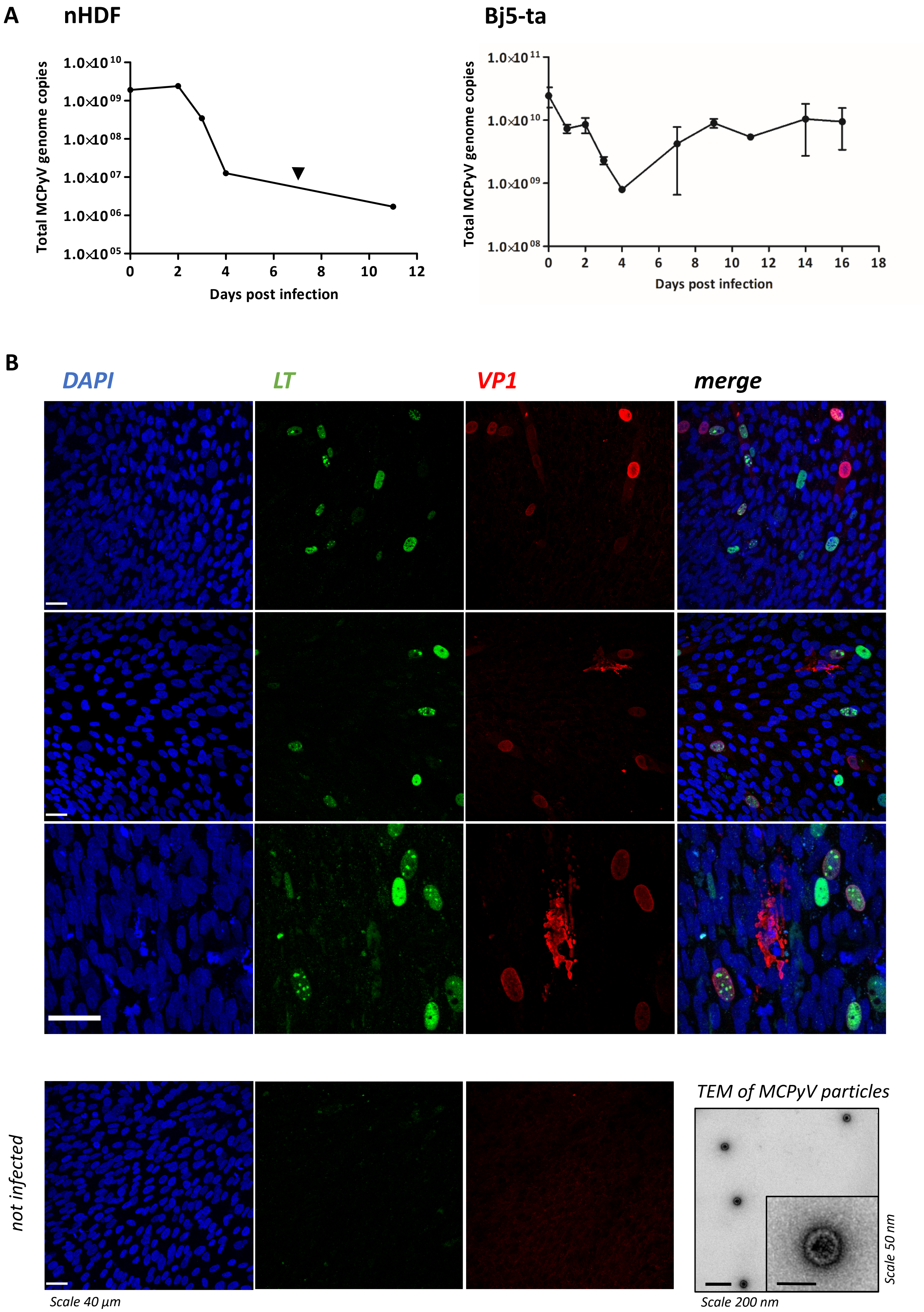

### supplementary Figure S2

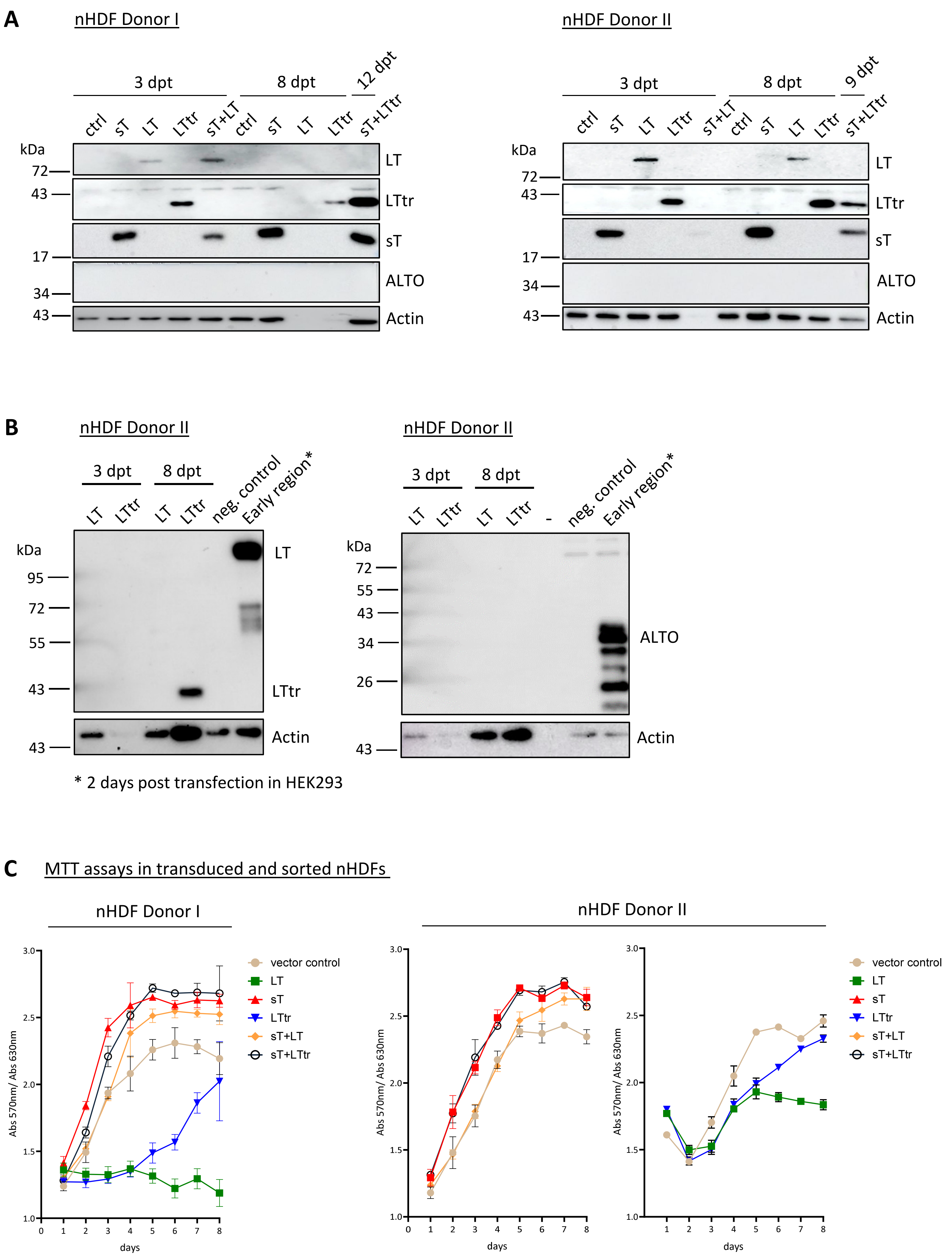

### supplementary Figure S3

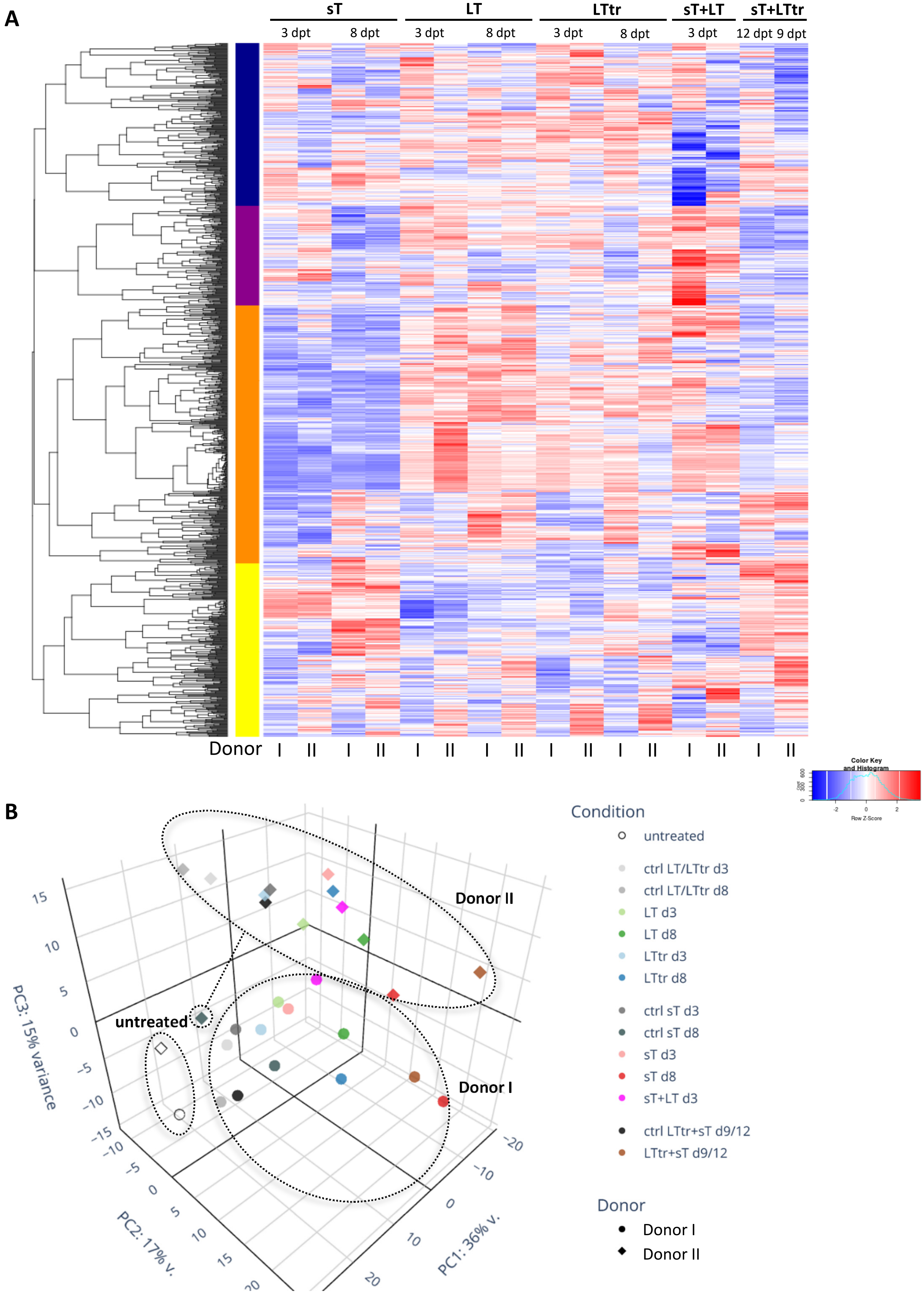

### supplementary Figure S5

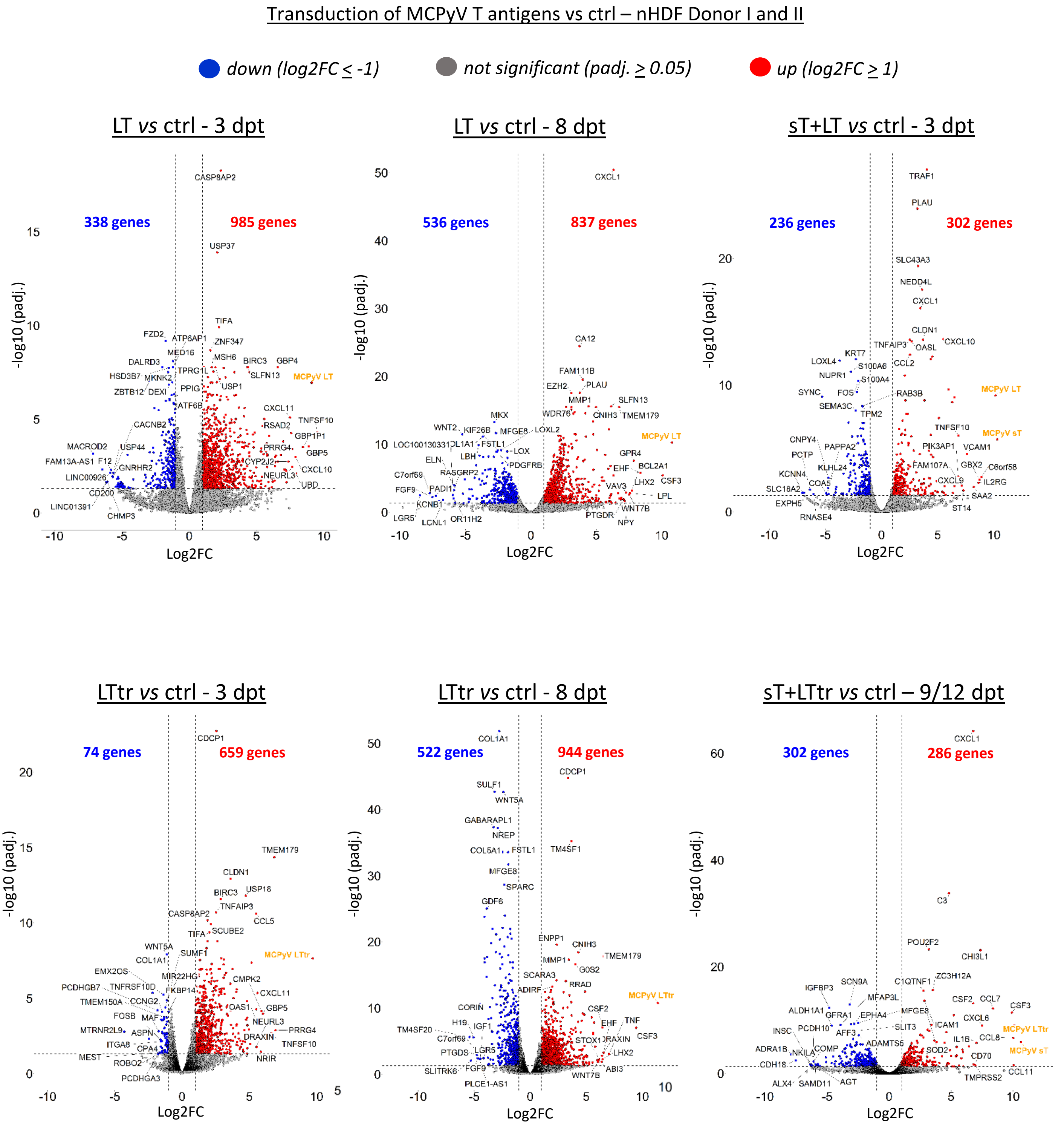

### supplementary Figure S6

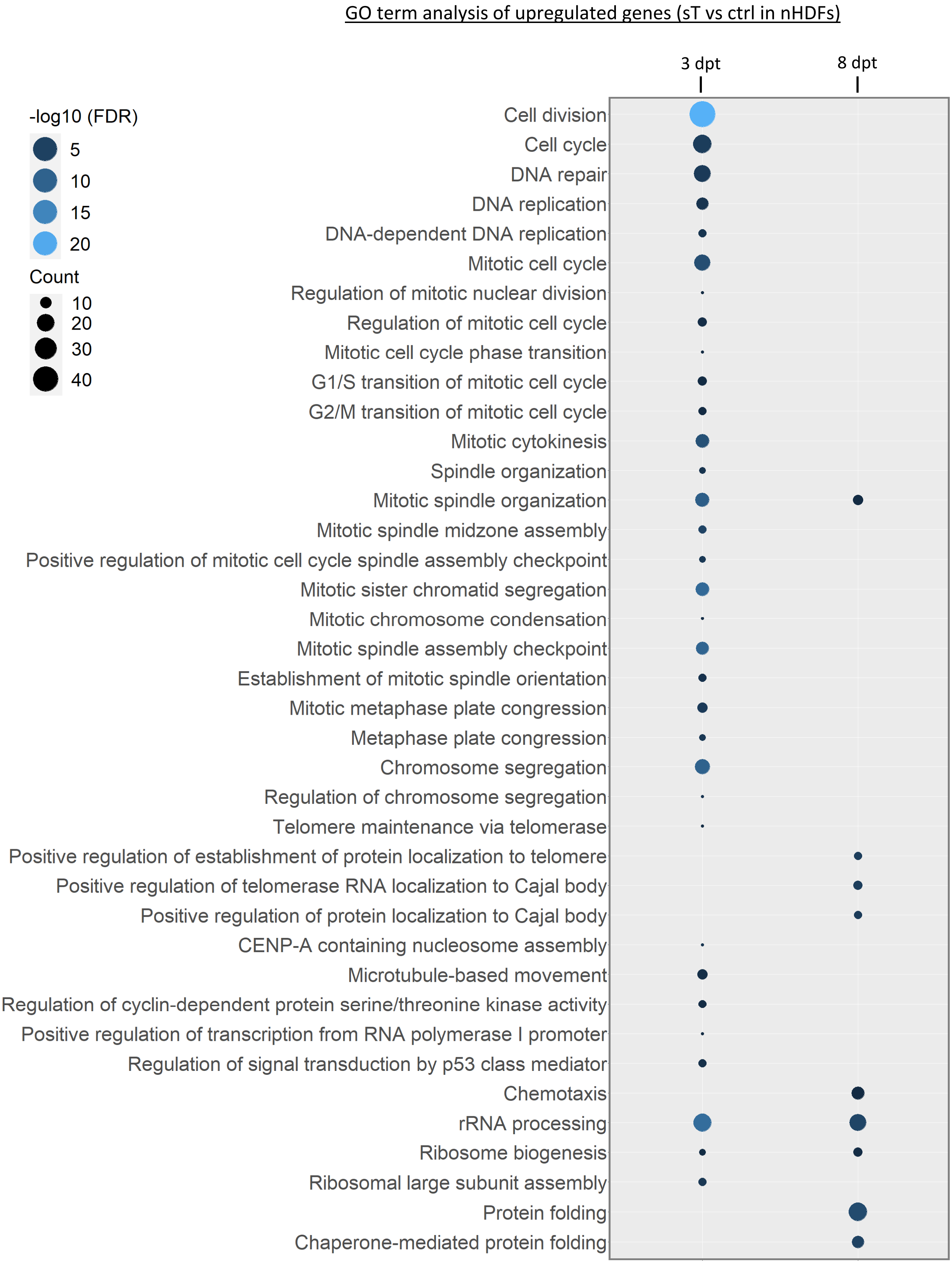

### supplementary Figure S7

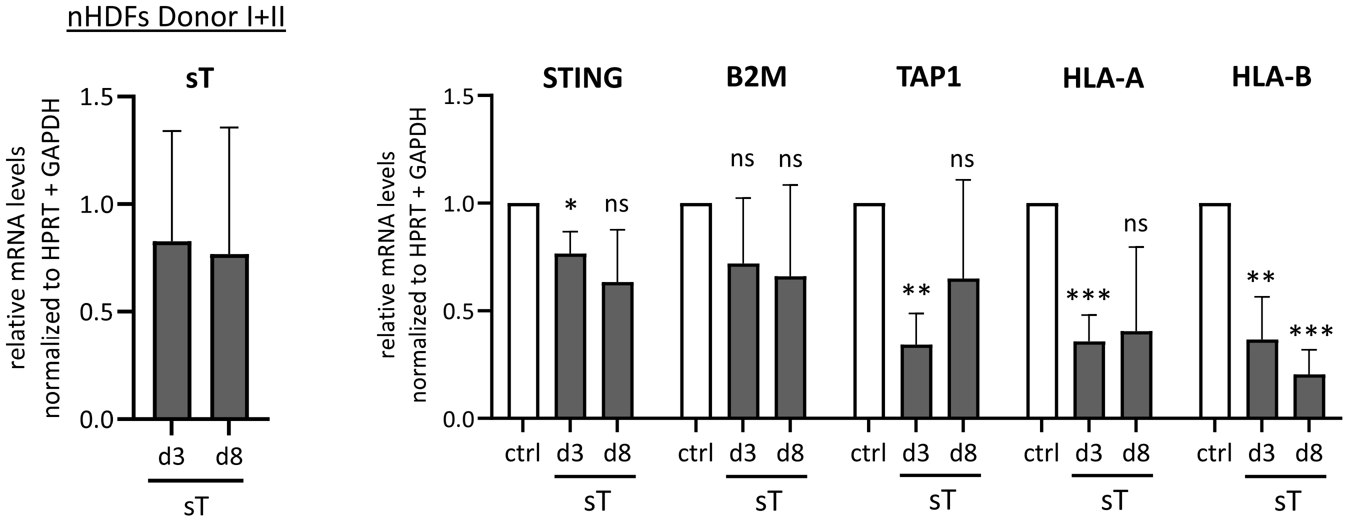

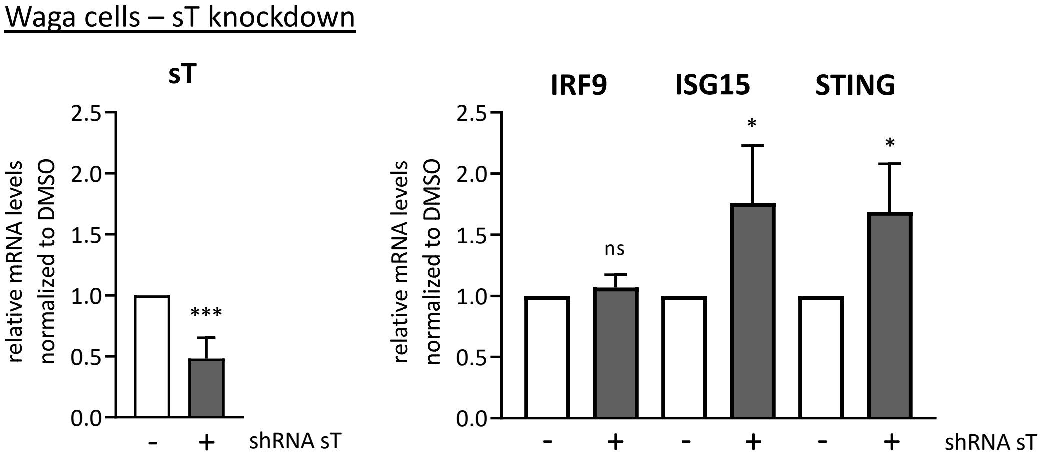

### supplementary Figure S8

**S8 Fig**

**A** Correlation of repressive histone marks and transcriptional changes (nHDFs)

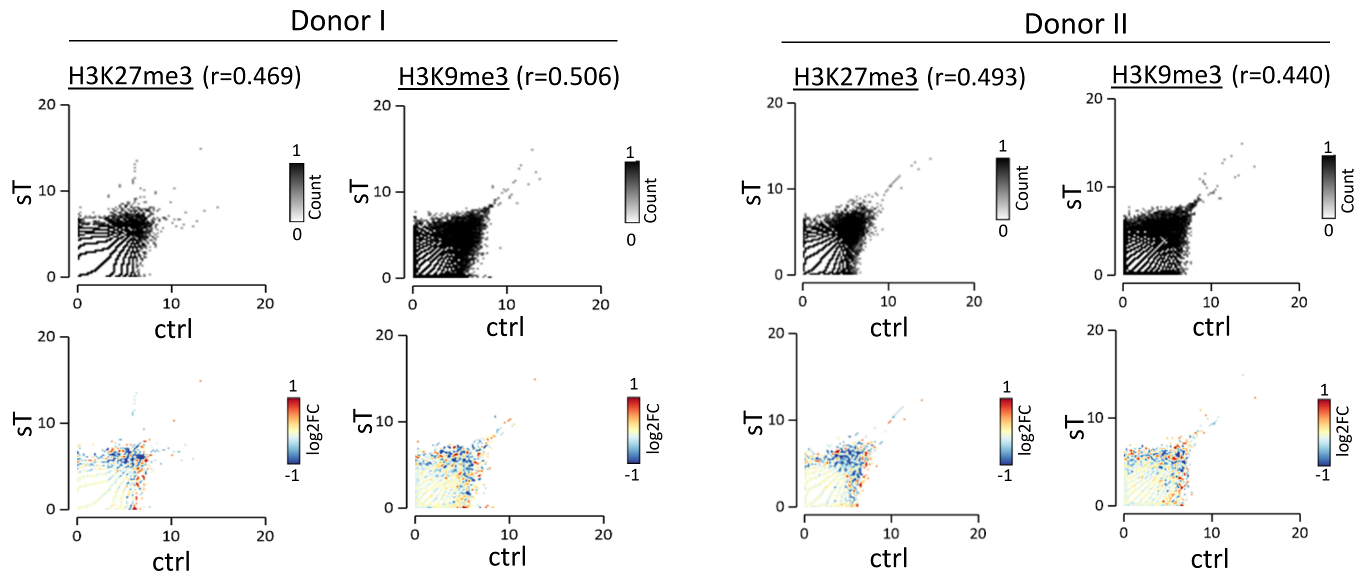


**B**

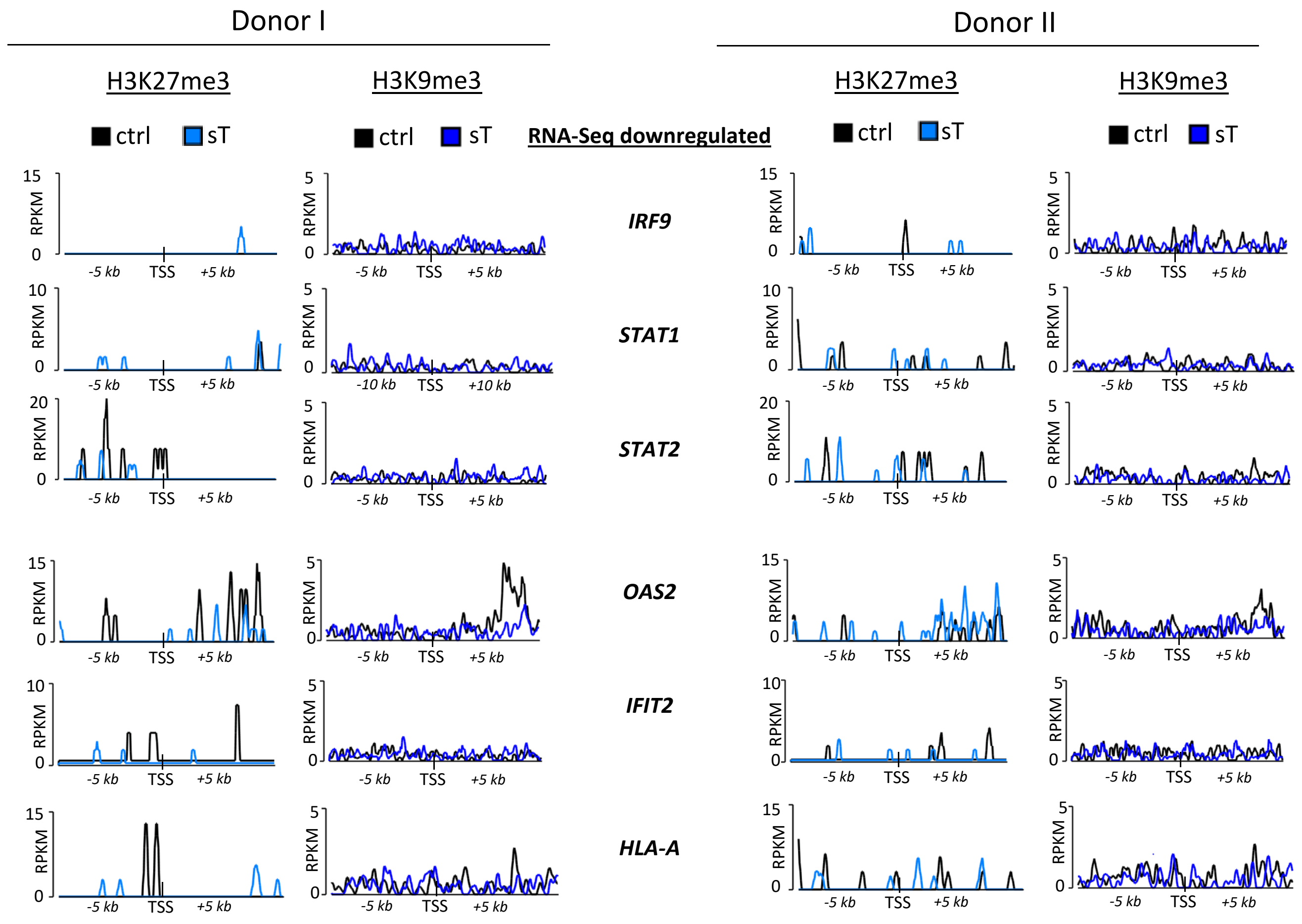

### supplementary Figure S9

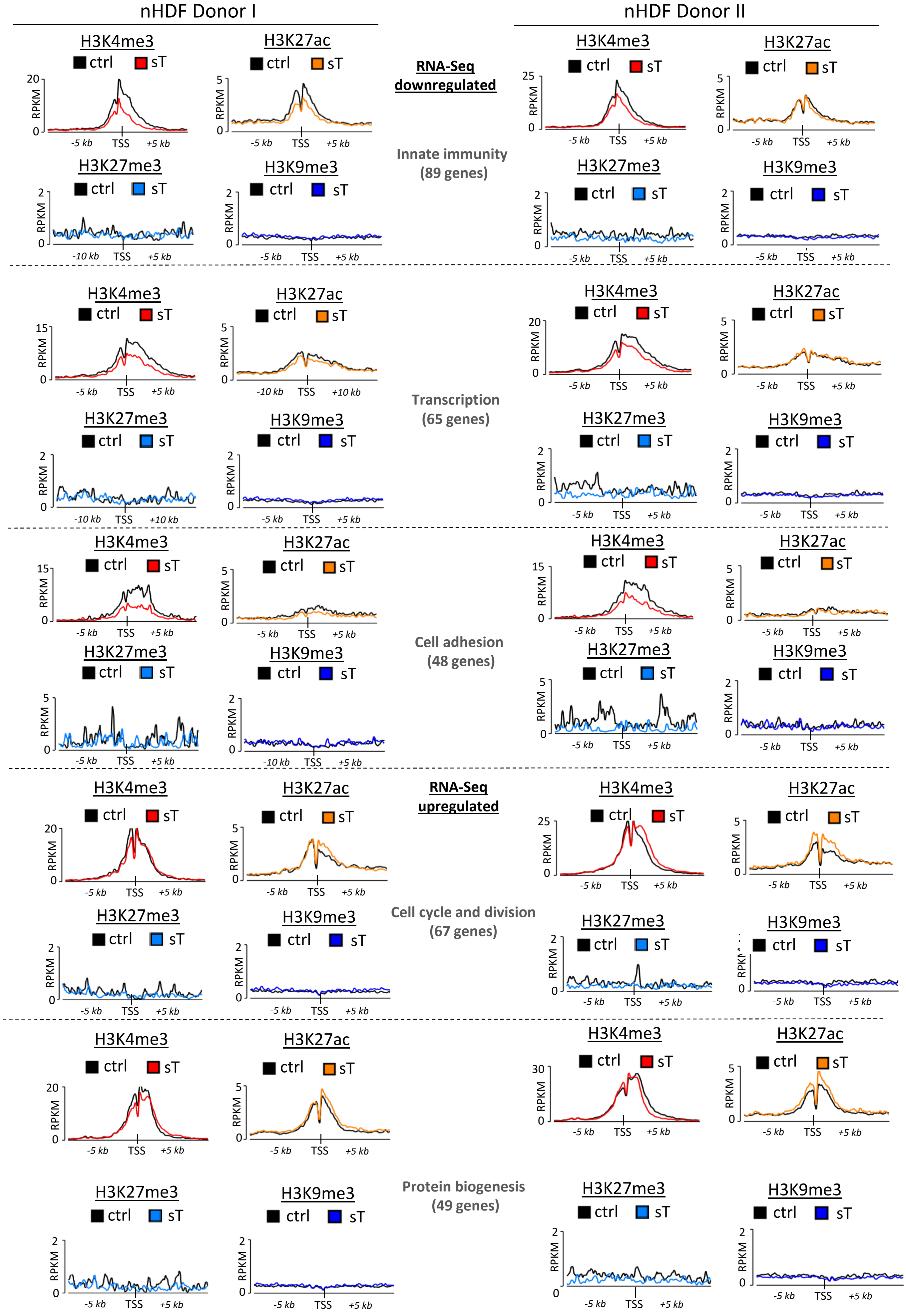

### supplementary Figure S10

#### S10 Fig

**A** IRF9 immunofluorescence analysis in nHDFs Donor I+II

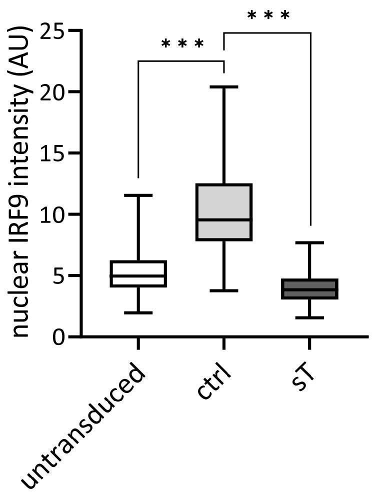


**B** nHDF Donor II

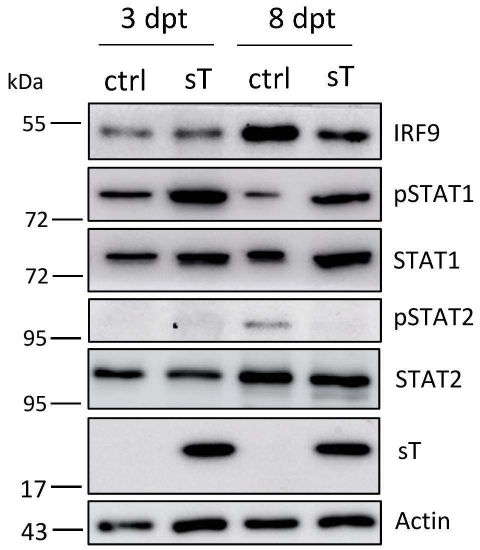

### supplementary Figure S11

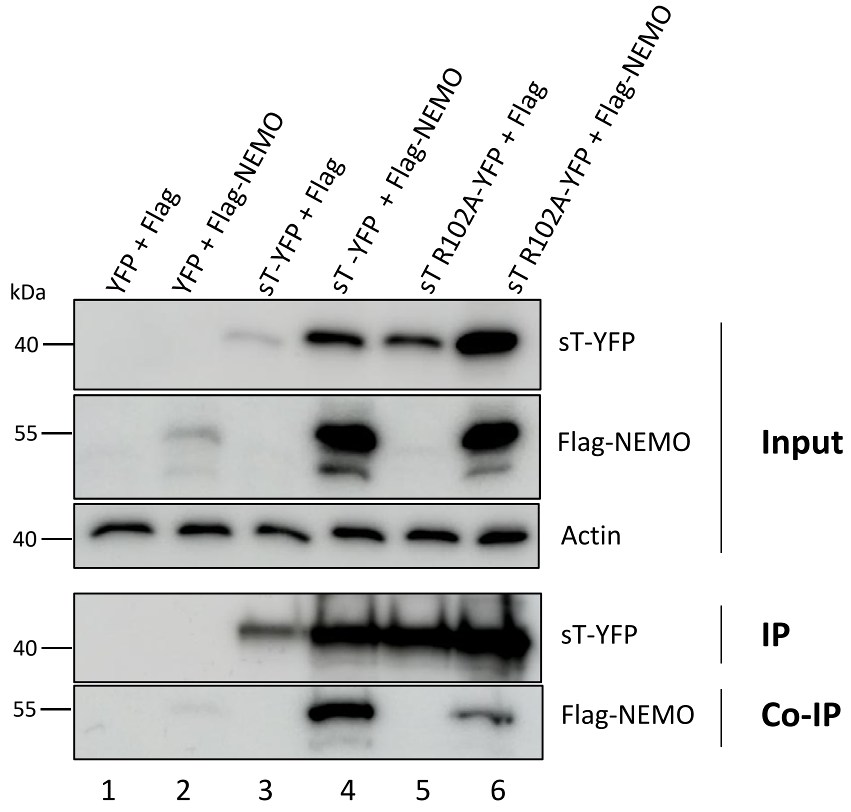

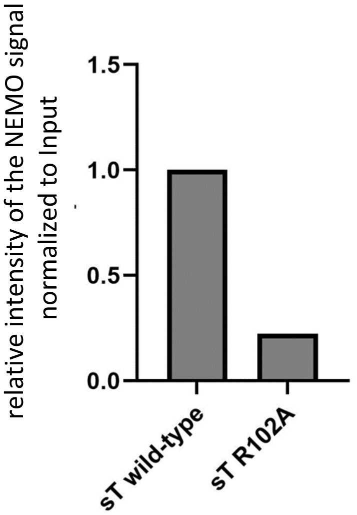

### supplementary Figure S13

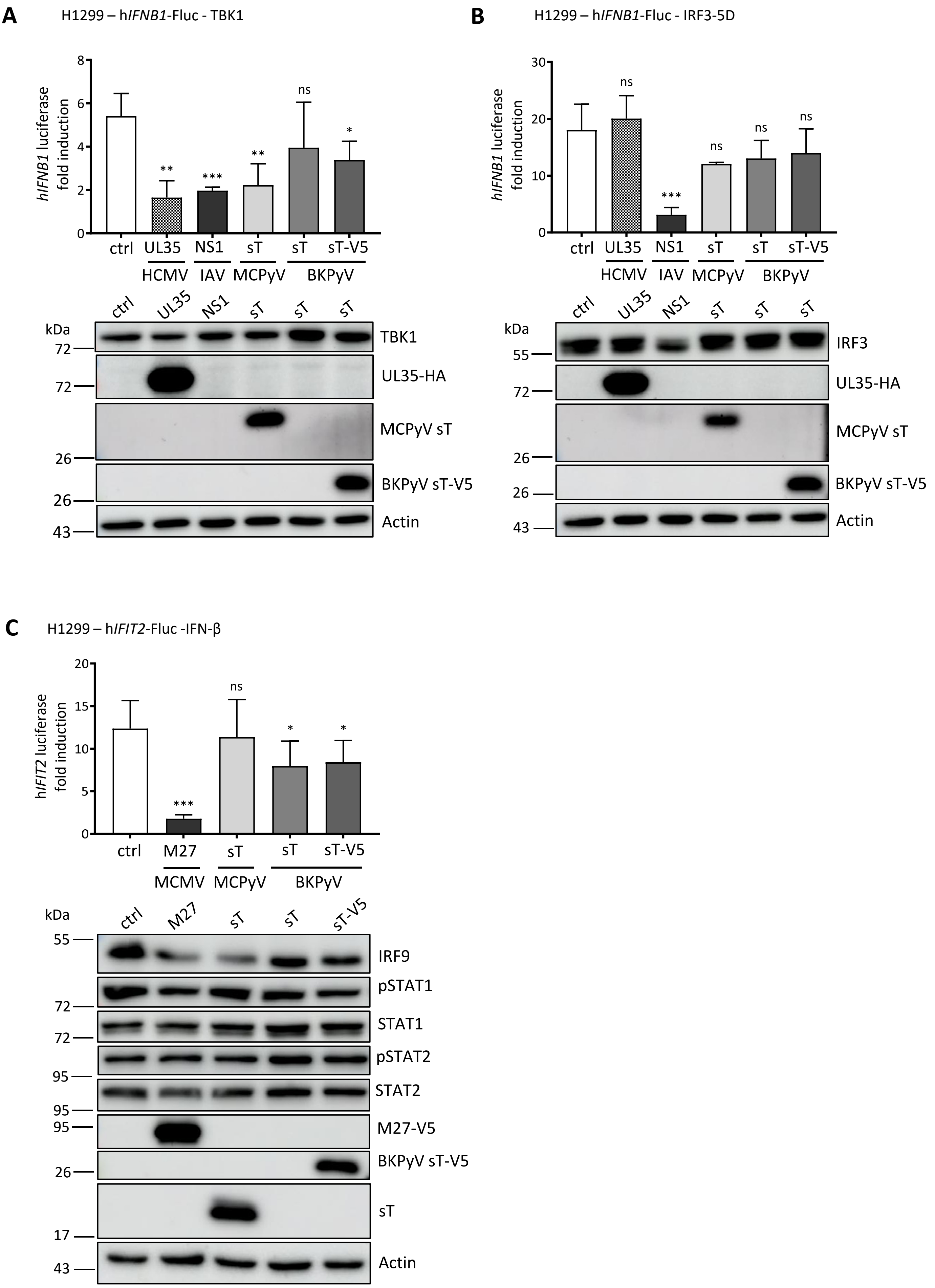

### supplementary Figure S14

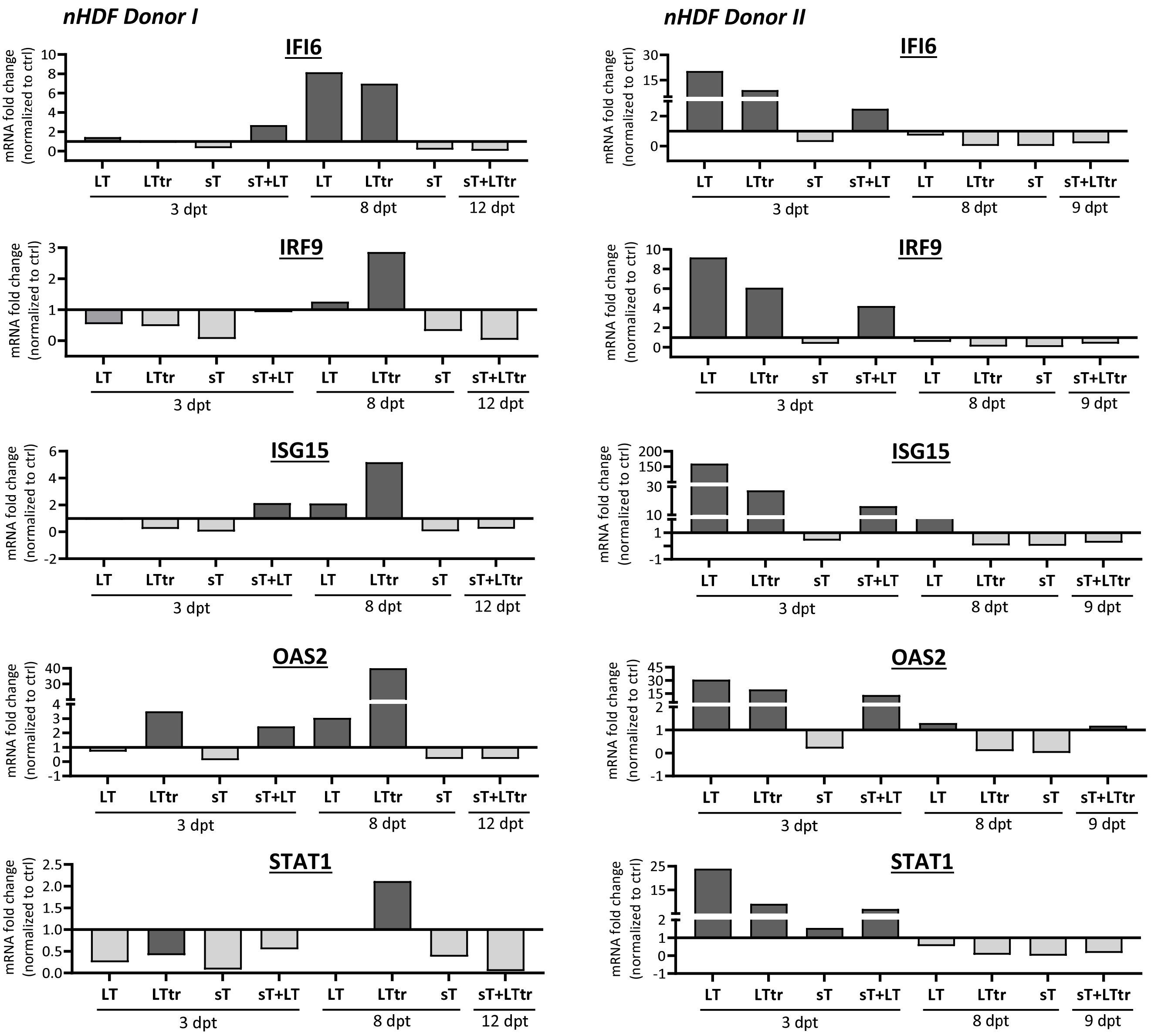
