## supplementary Figure S4 for "Merkel cell polyomavirus small tumor antigen contributes to immune evasion by interfering with type I interferon signaling"

### S4 Fig

#### Gene Ontology (Biological process) - cluster "BLUE"

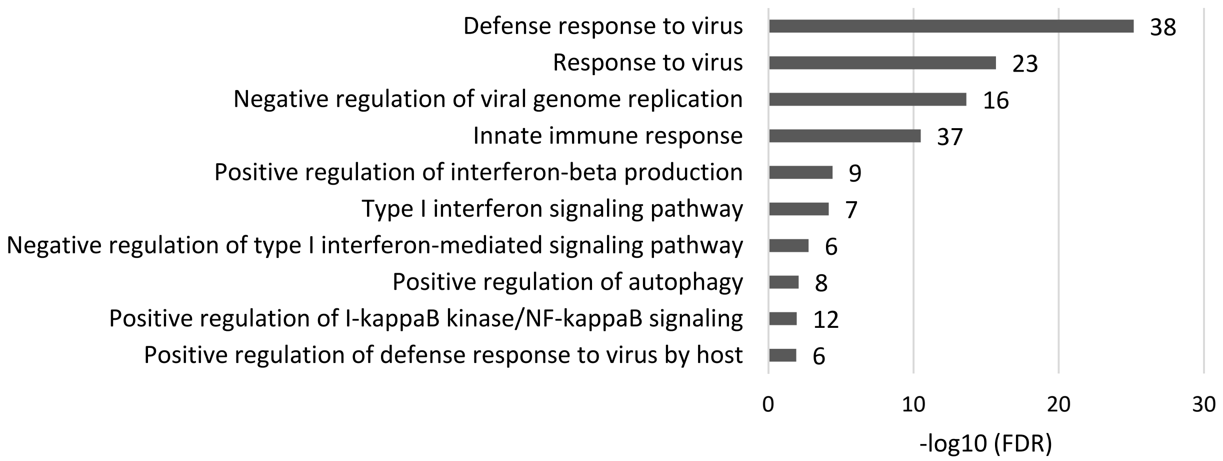


#### Gene Ontology (Biological process) - cluster "ORANGE"

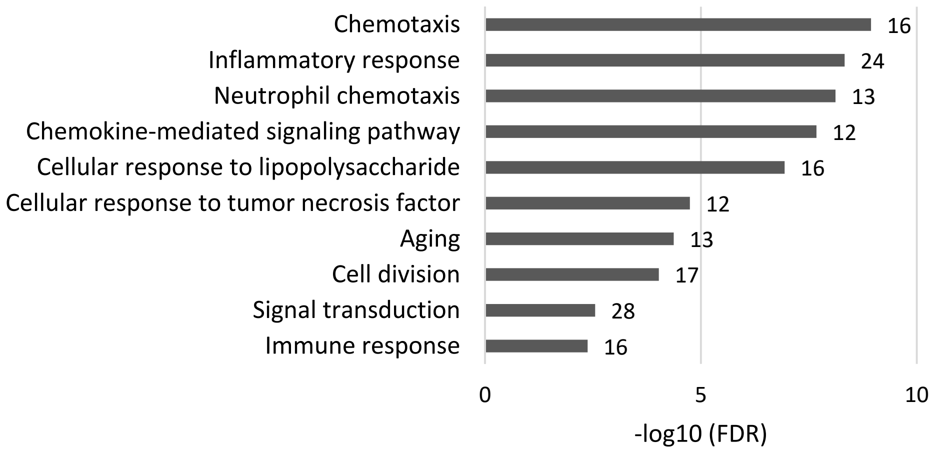


#### Gene Ontology (Biological process) - cluster "YELLOW"

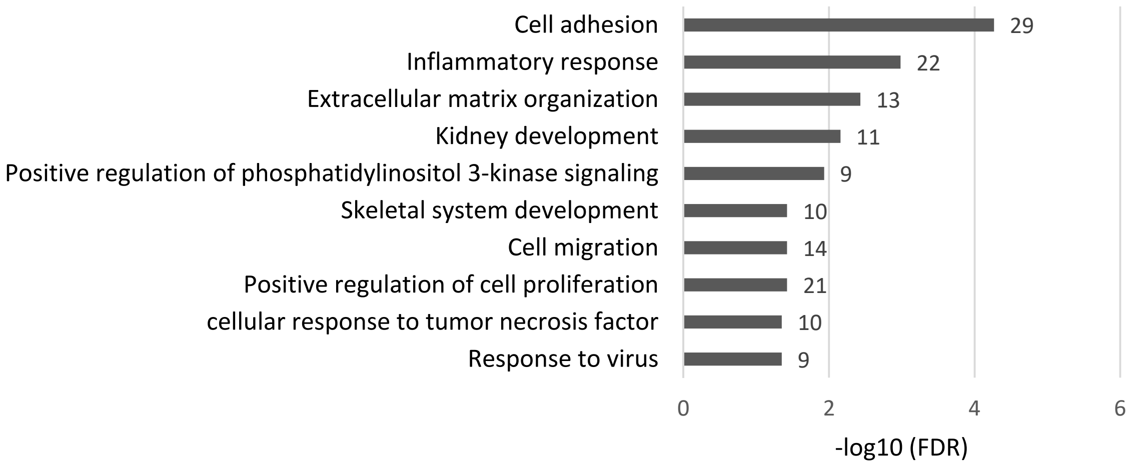
