## supplementary Figure S12 for "Merkel cell polyomavirus small tumor antigen contributes to immune evasion by interfering with type I interferon signaling"

### S12 Fig

**A** HEK293 – h*IFIT1*-Fluc -IFN-β immunoblots

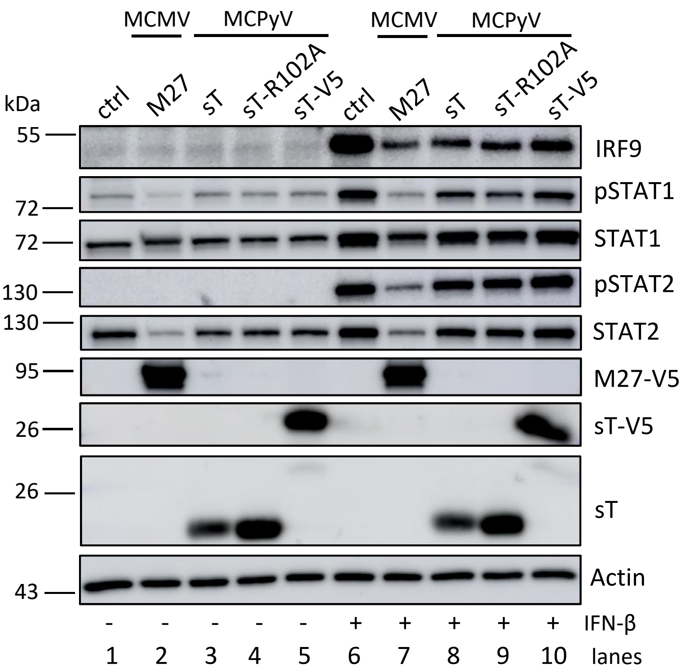


**B** HEK293 – h*IFIT2*-Fluc -IFN-β immunoblots

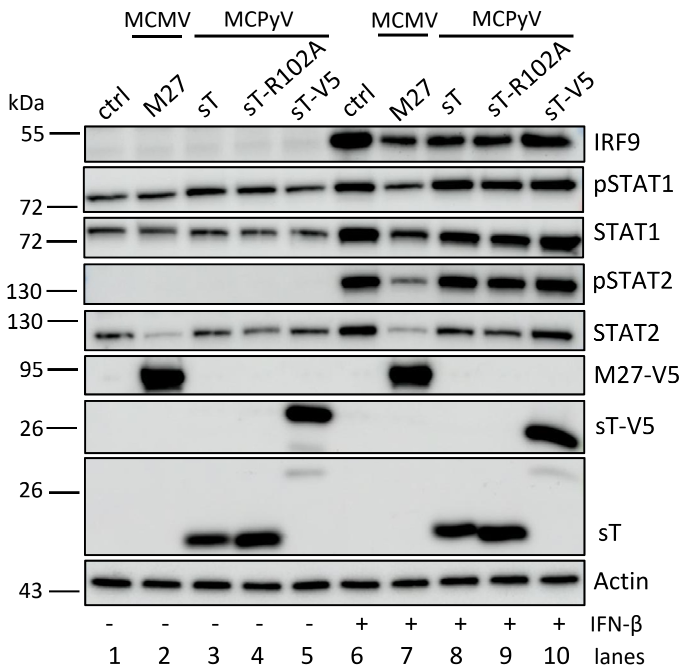


**C** HEK293 – m*MX1*-Fluc -IFN-β immunoblots

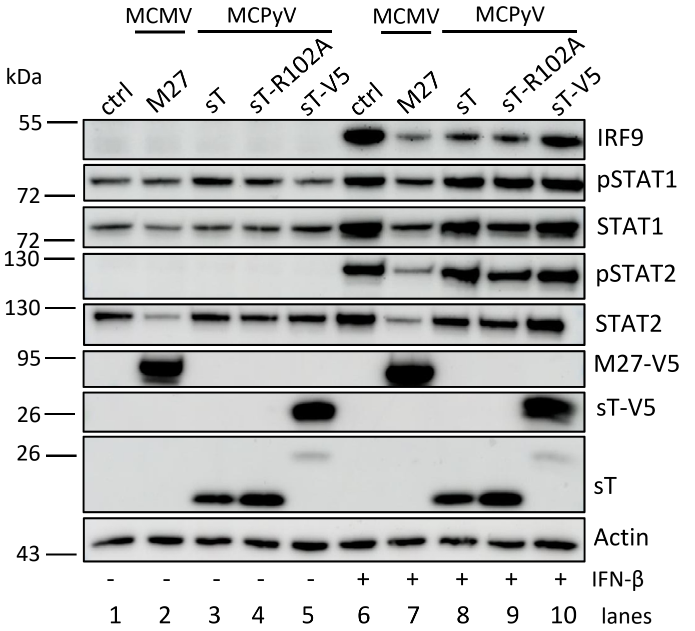
